# Dynamic gene expression analysis reveals distinct severity phases of immune and cellular dysregulation in COVID-19

**DOI:** 10.1101/2023.11.04.565404

**Authors:** Andy Y. An, Arjun Baghela, Peter Zhang, Travis M. Blimkie, Jeff Gauthier, Daniel E. Kaufmann, Erica Acton, Amy H.Y. Lee, Roger C. Levesque, Robert E.W. Hancock

## Abstract

**Background:** COVID-19 patients experience dynamic changes in immune and cellular function over time with potential clinical implications. However, there is insufficient research investigating, on a gene expression level, the mechanisms that become activated or suppressed over time as patients deteriorate or recover, which can inform use of repurposed and novel drugs as therapies.

**Objective:** To investigate longitudinal changes in gene expression profiles throughout the COVID-19 disease timeline.

**Methods:** Three-hundred whole blood samples from 128 adult patients were collected during hospitalization from COVID-19, with up to five samples per patient. Transcriptome sequencing (RNA-Seq), differential gene expression analysis and pathway enrichment was performed. Drug-gene set enrichment analysis was used to identify FDA-approved medications that could inhibit critical genes and proteins at each disease phase. Prognostic gene-expression signatures were generated using machine learning to distinguish 3 disease stages.

**Results:** Samples were longitudinally grouped by clinical criteria and gene expression into six disease phases: Mild, Moderate, Severe, Critical, Recovery, and Discharge. Distinct mechanisms with differing trajectories during COVID-19 hospitalization were apparent. Antiviral responses peaked early in COVID-19, while heme metabolism pathways became active much later during disease. Adaptive immune dysfunction, inflammation, and metabolic derangements were most pronounced during phases with higher disease severity, while hemostatic abnormalities were elevated early and persisted throughout the disease course. Drug-gene set enrichment analysis predicted repurposed medications for potential use, including platelet inhibitors in early disease, antidiabetic medications for patients with increased disease severity, and dasatinib throughout the disease course. Disease phases could be categorized using specific gene signatures for prognosis and treatment selection. Disease phases were also highly correlated to previously developed sepsis endotypes, indicating that severity and disease timing were significant contributors to heterogeneity observed in sepsis and COVID-19.

**Conclusions:** Higher temporal resolution of longitudinal mechanisms in COVID-19 revealed multiple immune and cellular changes that were activated at different phases of COVID-19. Understanding how a patient’s gene expression profile changes over time can permit more accurate risk stratification of patients and provide time-dependent personalized treatments with repurposed medications. This creates an opportunity for timely intervention before patients transition to a more severe phase, potentially accelerating patients to recovery.

## Introduction

As of October 2023, the COVID-19 pandemic has infected almost 700 million and killed 7-18 million people globally^1,2^. Patients with severe disease requiring hospitalization progress at different rates throughout their disease course. Individuals can manifest protracted, mild disease, or more severe disease including hospital stays for days to months, spending a portion of their stay in the intensive care unit (ICU), leading to high mortality rates of up to 32%^3^. In addition to the variation in disease duration, two patients in hospital for the same amount of time can have vastly different disease courses and immune profiles due to their variable progression rates. This reflects the reality of COVID-19 as a highly dynamic disease involving alternating and/or concurrent inflammation and immunosuppression^4,5^, which is a feature shared with sepsis (a life-threatening organ dysfunction due to an aberrant host response to infection^6^). Indeed, severe COVID-19 is clearly a form of viral sepsis^7,8^. Thus, in addition to demographic factors such as age, sex, ethnicity, and comorbidities and underlying mechanistic differences due to endotypes^9^, where individual patients are positioned on the COVID-19 disease timeline represents another source of patient heterogeneity with a substantial impact on appropriate patient management^10^.

Published longitudinal studies in COVID-19 to date have utilized small sample sizes leading to variable classifications of disease phases. One longitudinal transcriptomic study of peripheral blood mononuclear cells (PBMC) from nine patients classified patients into three severity stages and a recovery stage based on principal component analysis (PCA) of gene expression^4^. Another longitudinal PBMC study classified 18 patients into “treatment”, “convalescence”, and “rehabilitation” disease stages based on clinical symptoms^11^. A single-cell RNA-Seq study of 13 hospitalized patients classified patients into six “pseudo-times” ranging from “incremental” to “recovery/pre-discharge” based on disease severity^12^. Despite their major limitations of small sample size, these studies showed, during the most severe stages of COVID-19, consistent elevation of inflammatory mediators, dysregulated coagulation, and decreases in adaptive immunity, followed by reversal of these pathological processes during recovery stages.

Here we utilized clinical criteria and differential gene expression analysis to categorized patient samples into six distinct severity phases, using a large cohort of 128 hospitalized COVID-19 patients for which 300 samples were collected at different times in hospital. We identified shared and distinct underlying mechanistic processes occurring in each of these phases, leveraged these findings to identify potential phase-specific treatments, and then generated gene signatures that could classify patients into specific disease stages. Lastly, we explored how disease phase directly impacted on disease heterogeneity as reflected by endotype status.

## Methods

### Study design and patient recruitment

The Biobanque Québécoise de la COVID-19 (BQC-19; Quebec COVID-19 Biobank) recruited patients from ten hospitals across Quebec, Canada with confirmed SARS-CoV-2 infection^13^. This included 300 whole blood samples from 128 hospitalized patients (**Table 1**) that were collected during hospitalization, with individual patients having up to five samples collected in hospital. All patients were SARS-CoV-2 positive, adult (>18 years old), and hospitalized for COVID-19. Hospital samples were collected between April 2020 and February 2021, and due to this time frame, patients were infected by the ancestral strain, Alpha variant, or Beta variant and none were vaccinated prior to hospital admission^14^. In addition, 12 samples were also collected from 6 of these patients 3-7 months after discharge, all of whom reported no post-COVID symptoms during follow-up, to serve as controls.

**Table 1.**
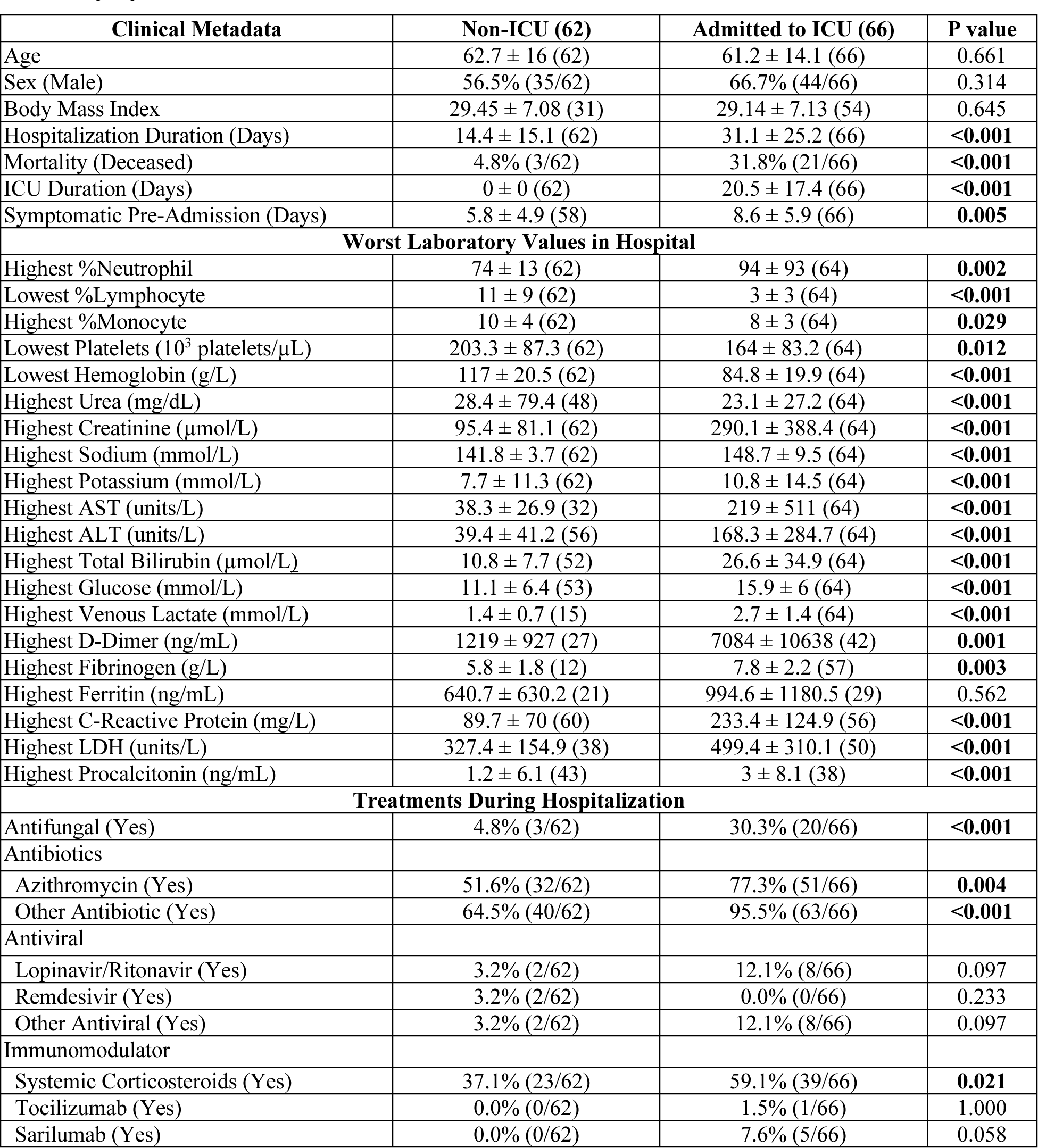
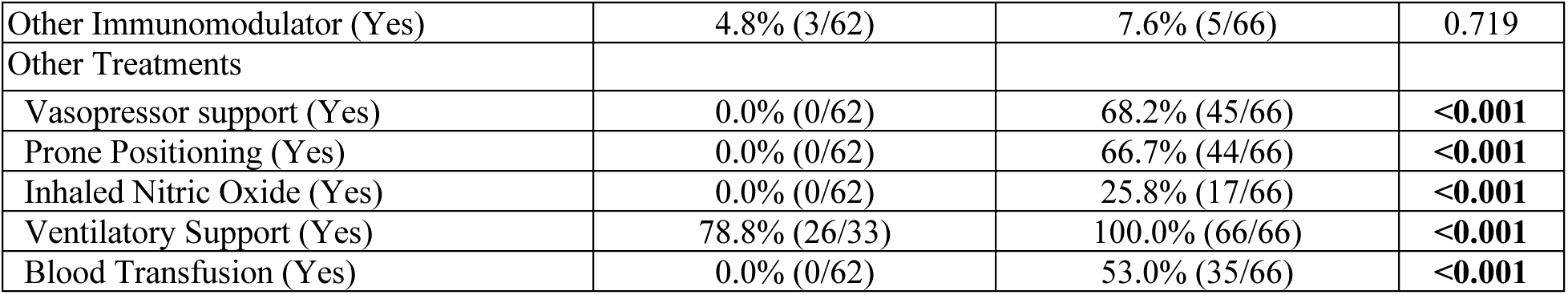
Demographics of patients in the BQC-19 biobank. For categorical variables, significance was tested using the Chi-squared test with Yates’s correction, or Fisher’s Exact test if any expected value was <5, and the percentage and fraction of patients fitting the category is displayed. For continuous variables, the Wilcoxon Rank-Sum test was used, and the mean ± standard deviation of the variable is displayed, with the number of patients assessed in brackets. Significant P-values (p <0.05) are bolded. Additional information including comorbidities and clinical symptoms can be found in **Table S4**.

| Clinical Metadata | Non-ICU (62) | Admitted to ICU (66) | P value |
| --- | --- | --- | --- |
| Age | 62.7 ± 16 (62) | 61.2 ± 14.1 (66) | 0.661 |
| Sex (Male) | 56.5% (35/62) | 66.7% (44/66) | 0.314 |
| Body Mass Index | 29.45 ± 7.08 (31) | 29.14 ± 7.13 (54) | 0.645 |
| Hospitalization Duration (Days) | 14.4 ± 15.1 (62) | 31.1 ± 25.2 (66) | **<0.001** |
| Mortality (Deceased) | 4.8% (3/62) | 31.8% (21/66) | **<0.001** |
| ICU Duration (Days) | 0 ± 0 (62) | 20.5 ± 17.4 (66) | **<0.001** |
| Symptomatic Pre-Admission (Days) | 5.8 ± 4.9 (58) | 8.6 ± 5.9 (66) | **0.005** |
| **Worst Laboratory Values in Hospital** |  |  |  |
| Highest %Neutrophil | 74 ± 13 (62) | 94 ± 93 (64) | **0.002** |
| Lowest %Lymphocyte | 11 ± 9 (62) | 3 ± 3 (64) | **<0.001** |
| Highest %Monocyte | 10 ± 4 (62) | 8 ± 3 (64) | **0.029** |
| Lowest Platelets ($10^3$ platelets/μL) | 203.3 ± 87.3 (62) | 164 ± 83.2 (64) | **0.012** |
| Lowest Hemoglobin (g/L) | 117 ± 20.5 (62) | 84.8 ± 19.9 (64) | **<0.001** |
| Highest Urea (mg/dL) | 28.4 ± 79.4 (48) | 23.1 ± 27.2 (64) | **<0.001** |
| Highest Creatinine (μmol/L) | 95.4 ± 81.1 (62) | 290.1 ± 388.4 (64) | **<0.001** |
| Highest Sodium (mmol/L) | 141.8 ± 3.7 (62) | 148.7 ± 9.5 (64) | **<0.001** |
| Highest Potassium (mmol/L) | 7.7 ± 11.3 (62) | 10.8 ± 14.5 (64) | **<0.001** |
| Highest AST (units/L) | 38.3 ± 26.9 (32) | 219 ± 511 (64) | **<0.001** |
| Highest ALT (units/L) | 39.4 ± 41.2 (56) | 168.3 ± 284.7 (64) | **<0.001** |
| Highest Total Bilirubin (μmol/L) | 10.8 ± 7.7 (52) | 26.6 ± 34.9 (64) | **<0.001** |
| Highest Glucose (mmol/L) | 11.1 ± 6.4 (53) | 15.9 ± 6 (64) | **<0.001** |
| Highest Venous Lactate (mmol/L) | 1.4 ± 0.7 (15) | 2.7 ± 1.4 (64) | **<0.001** |
| Highest D-Dimer (ng/mL) | 1219 ± 927 (27) | 7084 ± 10638 (42) | **0.001** |
| Highest Fibrinogen (g/L) | 5.8 ± 1.8 (12) | 7.8 ± 2.2 (57) | **0.003** |
| Highest Ferritin (ng/mL) | 640.7 ± 630.2 (21) | 994.6 ± 1180.5 (29) | 0.562 |
| Highest C-Reactive Protein (mg/L) | 89.7 ± 70 (60) | 233.4 ± 124.9 (56) | **<0.001** |
| Highest LDH (units/L) | 327.4 ± 154.9 (38) | 499.4 ± 310.1 (50) | **<0.001** |
| Highest Procalcitonin (ng/mL) | 1.2 ± 6.1 (43) | 3 ± 8.1 (38) | **<0.001** |
| **Treatments During Hospitalization** |  |  |  |
| Antifungal (Yes) | 4.8% (3/62) | 30.3% (20/66) | **<0.001** |
| Antibiotics |  |  |  |
| Azithromycin (Yes) | 51.6% (32/62) | 77.3% (51/66) | **0.004** |
| Other Antibiotic (Yes) | 64.5% (40/62) | 95.5% (63/66) | **<0.001** |
| Antiviral |  |  |  |
| Lopinavir/Ritonavir (Yes) | 3.2% (2/62) | 12.1% (8/66) | 0.097 |
| Remdesivir (Yes) | 3.2% (2/62) | 0.0% (0/66) | 0.233 |
| Other Antiviral (Yes) | 3.2% (2/62) | 12.1% (8/66) | 0.097 |
| Immunomodulator |  |  |  |
| Systemic Corticosteroids (Yes) | 37.1% (23/62) | 59.1% (39/66) | **0.021** |
| Tocilizumab (Yes) | 0.0% (0/62) | 1.5% (1/66) | 1.000 |
| Sarilumab (Yes) | 0.0% (0/62) | 7.6% (5/66) | 0.058 |
| Other Immunomodulator (Yes) | 4.8% (3/62) | 7.6% (5/66) | 0.719 |
| Other Treatments |  |  |  |
| Vasopressor support (Yes) | 0.0% (0/62) | 68.2% (45/66) | <0.001 |
| Prone Positioning (Yes) | 0.0% (0/62) | 66.7% (44/66) | <0.001 |
| Inhaled Nitric Oxide (Yes) | 0.0% (0/62) | 25.8% (17/66) | <0.001 |
| Ventilatory Support (Yes) | 78.8% (26/33) | 100.0% (66/66) | <0.001 |
| Blood Transfusion (Yes) | 0.0% (0/62) | 53.0% (35/66) | <0.001 |

### Classification of samples into disease phases

Samples collected at hospitalization were classified into one of six disease phases to simulate the various phases that patients transition through during disease, considering both Sequential Organ Failure Assessment (SOFA) score and whether patients were hospitalized in the ICU (**Figure S1**). “Mild” samples were non-ICU samples with a SOFA score <2. “Moderate” samples were non-ICU samples that had SOFA scores between 2 and 6. “Severe” samples were from patients in the ICU with SOFA scores between 2 and 11, or non-ICU samples with SOFA scores >6. “Critical” samples were ICU samples from patients with SOFA scores ≥12. “Recovery” samples were ICU samples that were part of a downward trajectory in SOFA score (*i.e.*, the samples collected before and after had higher and lower SOFA scores, respectively, or the patient was discharged from the ICU soon after sample collection). “Discharge” samples were non-ICU samples collected after ICU discharge, were part of a downward trajectory in SOFA score, or had a SOFA score <2 and were collected in the second half of hospitalization duration. SOFA score cut-offs of 2, 6, and 12 were used based on mortality rates from a multicenter study on SOFA scores in the ICU^15^: a SOFA score <2 does not satisfy Sepsis-3 criteria^6^, SOFA scores of 2 to 6 are associated with a mortality rate of <10%, SOFA scores 7 to 11 with a mortality rate of 15-45%, and SOFA scores ≥12 with a mortality rate of >50%. In total, there were 19 Mild, 53 Moderate, 119 Severe, 26 Critical, 22 Recovery, and 61 Discharge samples. The classified phases also trended similarly when using the World Health Organization COVID-19 Clinical Progression Scale^16^ instead of SOFA score (**Figure S2**). These phases were also significantly different in various clinical metadata which trended similarly to SOFA score (*i.e.*, peaking at the Critical phase) (**Table S1**), including neutrophil counts, urea, creatinine, aspartate transaminase (AST), C-reactive protein, and procalcitonin. Other clinical metadata trended in the opposite direction (*i.e.*, lowest at the Critical phase), including lymphocyte and monocyte counts, platelets, diastolic blood pressure, hemoglobin, and albumin. Phases were also later paired into 3 disease *stages*, namely Initial (Mild, Moderate), Peak (Severe, Critical), and Convalescence (Recovery, Discharge) to obtain sufficient samples to enable identification of diagnostic gene-expression signatures specific to each stage.

### RNA-Seq and alignment

Approximately 2.5 ml of whole blood was collected for RNA-Seq. Blood was collected into PAXgene Blood RNA tubes (BD Biosciences; San Jose, CA, USA) to ensure stabilization of intracellular RNA. Total RNA was extracted using the PAXgene Blood RNA Kit (Qiagen; Germantown, MD, USA). Quantification and quality measures for total RNA were obtained using a LabChip GXII instrument (Waltham, MA, USA). Poly-adenylated RNA was captured using NEBNext Poly(A) mRNA Magnetic Isolation Module (NEB; Ipswich, USA) and cDNA libraries were prepared using the NEBNext RNA First Strand Synthesis Module, NEBNext Ultra Directional RNA Second Strand Synthesis Module, and NEBNext Ultra II DNA Library Prep Kit for Illumina (NEB; Ipswich, USA). RNA-Seq was then performed at a depth of 50M reads/sample on an Illumina NovaSeq 6000 S4 instrument of 100 base-pair long paired-end sequence reads (excluding adapter/index sequences). All bioinformatic analyses were performed in the programming language R (v4.2.2)^17^. A standard RNA-Seq processing pipeline was followed, including *FastQC* (v0.11.9)^18^ and *MultiQC* (v1.6)^19^ quality control, *STAR* (v2.7.9a)^20^ alignment to the human genome (Ensembl GRCh38.104), and assessing read counts using *HTSeq-count* (v0.11.3)^21^. All samples analyzed had more than one million total reads.

### Differential gene expression and pathway analysis

The count matrix was pre-filtered to remove globin genes (*HBA1*, *HBA2*, *HBB*, *HBD, HBG1, HBG2*) and low-count genes (mean counts across all samples <10) prior to differential expression analysis, resulting in a gene universe of 18,562 ENSEMBL gene IDs. The package *DESeq2*^22^ was used to identify differentially expressed (DE) genes. Sequencing batch and sex were modelled as covariates in the *DESeq2* model, and the Wald test was used for hypothesis testing. Comparisons between phases/stages to each other and to controls (follow-up samples) were performed to identify differentially expressed (DE) genes. To generate PCA plots, the *PCAtools* package^23^ was used, removing the bottom 10% of genes with low variance. To understand underlying pathophysiology, pathway enrichment using DE genes was performed using the *SIGORA* package (v3.1.1)^24^ with the Reactome pathway database^25^. Reactome pathways were considered significantly enriched with an adjusted p-value <0.001 (Bonferroni multiple test correction) as was recommended in *SIGORA*. To supplement and validate *SIGORA* pathway enrichment results, the Hallmark gene sets from the Molecular Signatures Database (MSigDB) were also analyzed^26^. Gene sets were considered significantly enriched with an adjusted p-value <0.05 (Benjamini-Hochberg multiple test correction) and q-value <0.2, based on the default settings of the *enricher* function in the package *clusterProfiler* (v4.2.2)^27^. Enrichment was performed separately on up- and down-regulated DE genes. Pathways and gene sets were considered “upregulated” if the genes in these pathways or gene sets were overrepresented in upregulated DE genes when compared to their prevalence in the gene universe, suggesting an increase in their function or activity, and vice versa for “downregulated”. Gene set variation analysis (GSVA) was performed using *DESeq2* variance-stabilized transformed counts with the *GSVA* package^28^ to identify enrichment scores of Hallmark gene sets of interest.

### Disease stage signature generation

The top 50 upregulated DE genes with the highest fold changes from each disease stage (Initial, Peak, and Convalescence) when compared to all other phases (*e.g.*, Convalescent vs. Initial and Peak) were chosen as a preliminary gene signature for each stage. To create a condensed gene signature of each stage, least absolute shrinkage and selection operator (LASSO) regression from the package *glmnet* (v4.1.6)^29^ was used to reduce the number of genes in the signature, with the disease stage as the response variable and the preliminary gene expression signature as predictor variables. A ten-fold cross-validation was performed to find the best lambda value that produced the lowest test mean squared error, which was then used to develop the multinomial LASSO regression model. Genes with coefficients shrunk to zero were removed, creating a condensed gene signature. GSVA was performed using *DESeq2* variance-stabilized transformed counts with the *GSVA* package^28^ to classify samples into disease stages based on the preliminary and condensed gene signatures. The accuracy of the classification was calculated by the sum of true positives and true negatives, divided by the total number of samples.

## Results

### Distinct immune and cellular pathways occur at different COVID-19 disease phases

To identify the mechanistic progression of COVID-19, differential gene expression was performed to identify DE genes in each disease phase (**Figure S3**). The validity of the disease phases was revealed by the high total number of DE genes in each phase with 2,003 in Mild, increasing to 2,220 in Moderate and 5,867 in Severe, then peaking at 8,542 DE genes in Critical, after which numbers decreased to 3,556 DE genes in Recovery and 1,202 DE genes in Discharge. This indicated increasing dysregulation compared to controls as patients became critically ill and the reverse when patients began to recover. Interestingly, 734 shared genes were DE at all phases, suggesting that certain processes remained dysregulated throughout the disease timeline, even when patients were ready to be discharged. PCA on these samples showed that samples clustered primarily based on the severity of their disease phase, with Severe and Critical clustering together and the remaining less-severe phases generally clustering with controls (**Figure S4A**); this was consistent with the potential for higher-level groupings into stages (**Figure S4B**).

Pathway enrichment was performed on phase-specific DE genes to identify processes that were dysregulated at each phase (**Figure 1**). Notably, a variety of immune pathways were highly dysregulated, albeit at different phases (**Figure 1A**). “Interferon α/β Signaling” and “Interferon Signaling” were highly enriched by upregulated genes from Mild to Critical phases but were no longer enriched in the Recovery and Discharge phases, consistent with an early antiviral response that diminished as patients recovered; this was recapitulated by the Hallmark gene sets for “Interferon-α Response” and “Interferon-γ Response” (**Figure 1B**). Conversely, the adaptive immune pathway “Immunoregulatory Interactions”, the signaling pathway “DAP12 Signaling” (involved in NK cells and myeloid cell activation)^30^, and the T cell signaling gene set “IL2 STAT5 Signaling” were only highly enriched by downregulated genes from Moderate to Recovery phases consistent with immunosuppression in these phases. Other adaptive immune pathways were also downregulated in later phases, including “Co-stimulation by the CD28 Family”, “MHC class II Antigen Presentation”, and the “Allograft Rejection” gene set. Immune pathways that were enriched by upregulated genes only in the high severity phases included interleukin signaling pathways such as “IL-1 Signaling” and “IL-4/13 Signaling”, “ER-Phagosome Pathway”, and “Chemokine Receptors Bind Chemokines”. Conversely, “Neutrophil Degranulation” was significantly upregulated at all phases, suggesting underlying processes that were activated throughout the entire disease course up to discharge. The enrichment of a greater number of immune pathways in the worst-disease phases indicated that immune dysfunction is a hallmark of severe but not mild COVID-19, as further supported by the peaking, during the Critical phase, of these pathways (revealed by median log_2_ fold change of the constituent genes) (**Figure 1C**).

**Figure 1.**
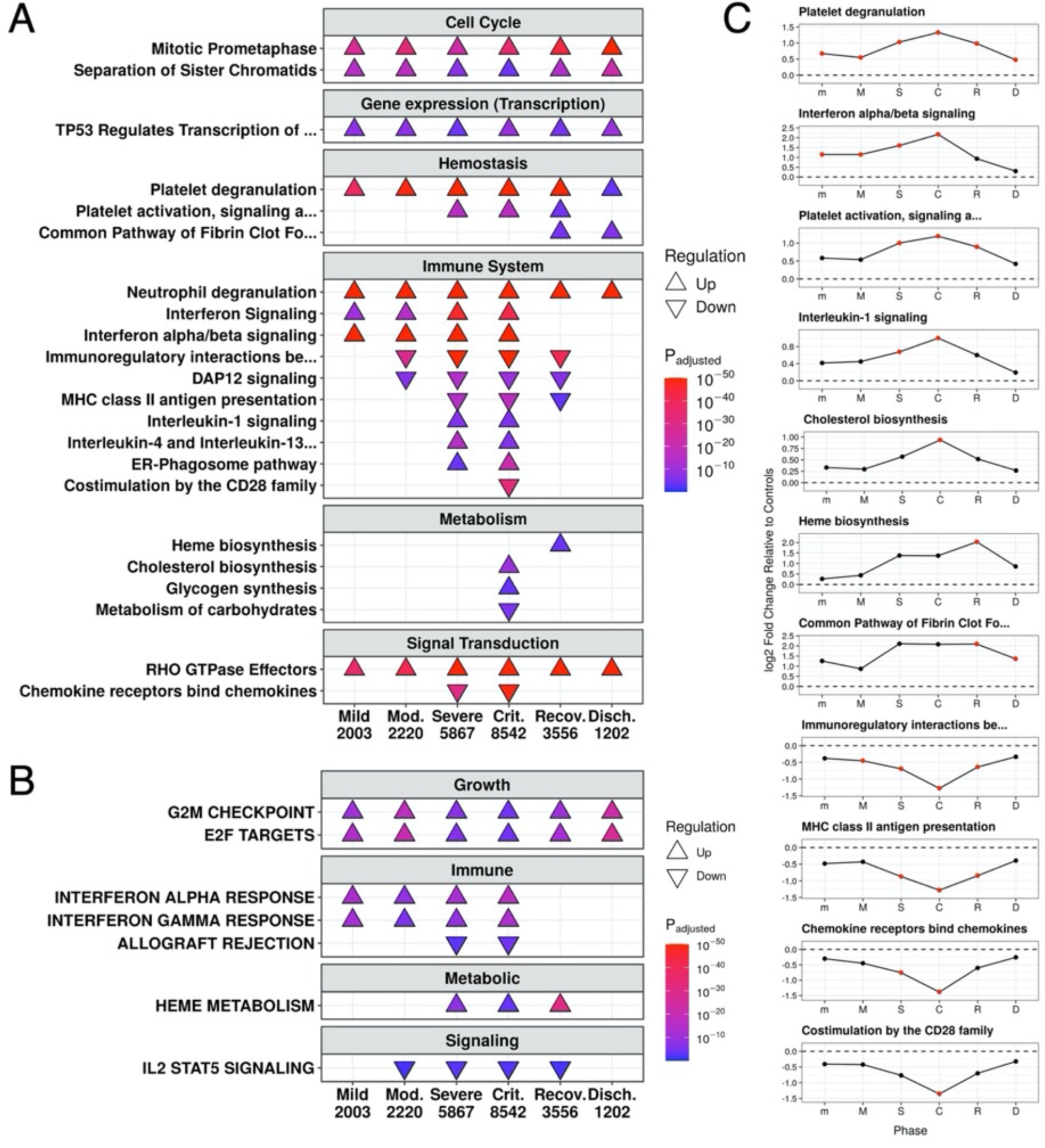
Mechanistic changes occur as patients progress through different disease phases. **A:** Subset of enriched Reactome pathways, all enriched pathways shown in **Figure S5**. **B:** Subset of enriched Hallmark gene sets, with all enriched gene sets shown in **Figure S6**. The total number of DE genes in each comparison are shown under the label. **C:** Line graphs representing trends of pathway enrichment in **A**, phases are abbreviated as their first letter except m = mild. Points indicate the median log_2_ fold change of the set of genes of each pathway that were DE at any of the phases. Red points indicate significant enrichment of pathway at that phase, as indicated in **A**.

Various other cellular processes were also dysregulated over time. Multiple cell cycle pathways such as “Mitotic Prometaphase” were highly enriched by upregulated genes throughout the entire disease process, as were related pathways such as “TP53 Regulates Transcription of Genes involved in G1 Cell Cycle Arrest” and “RHO GTPase Effectors”. These play a role in cell-cycle progression^31^, as do the Hallmark gene sets “G2M Checkpoint” and “E2F Targets” that exhibited similar patterns of enrichment. “Platelet degranulation” was another pathway that was upregulated throughout the disease process, although the “Platelet activation, signaling, and aggregation” pathway was only upregulated from the Severe to Recovery phases. The “Common Pathway of Fibrin Clot Formation” was also upregulated only in the Recovery and Discharge phases, consistent with lingering hemostatic dysfunction up to discharge. Late activation of heme metabolism in COVID-19 was also seen in this cohort. The “Heme Biosynthesis” pathway was upregulated only in the Recovery phase, while the “Heme Metabolism” gene set was only upregulated from the Severe to Recovery phases. A wide variety of metabolic pathways were enriched only during the Critical phase, including “Cholesterol Biosynthesis”, “Metabolism of Carbohydrates”, and “Glycogen Synthesis”, highlighting metabolic derangements in the most critical period of sickness. To further confirm these trends, GSVA enrichment was performed using genes in each pathway, and similar trends were observed including an early interferon response, a late heme metabolic response, and IL-1 signaling and platelet degranulation processes peaking in the Critical phase (**Figure S7**).

Since a proportion of patients in this cohort died in hospital, we then determined whether these patients differed in their trajectories. Of the 64 samples collected from patients who eventually died, 59 were classified as either Severe (45) or Critical (14). These were then compared to Severe and Critical phase samples in patients who ultimately recovered and were discharged from the hospital. Most of the differences occurred during the Critical phase, with 1,696 DE genes (**Figure S8**). Comparing non-survivors to survivors during this phase, there was upregulation of “Neutrophil Degranulation” and downregulation of “Immunoregulatory Interactions” and various hemostasis-related pathways, consistent with immunosuppression, aberrant inflammation, and decreased platelet function, despite patients having similar clinical presentation (**Table S2**) and severity (total SOFA scores and WHO severity scale values were not significantly different). The only observed difference in clinical metadata between these Critical samples of non-survivors and survivors was significantly lower platelet counts in patients who died (and a corresponding higher coagulation SOFA component score), corresponding to the hemostasis differences observed through pathway enrichment. Overall, these results highlighted disease mechanisms with varying trajectories in COVID-19.

### Repurposed drugs have potential uses for phase-specific treatment

Based on the multiple phase-dependent processes identified, drug-gene interaction analysis was used to identify repurposed drugs that can potentially inhibit critical mechanisms that are dysfunctional. The Drug Signatures Database (DSigDB)^32^ is a database of drug-gene interactions of FDA-approved medications determined through *in vitro* cell culture studies. By applying this database to phase-specific upregulated genes, approved drugs that have the potential to target each phase or multiple phases were identified. Multiple drugs unique to specific phases were enriched (**Figure 2**). Unique drugs with signatures that were only enriched in the “Mild” and “Moderate” phases (“Initial” stage) included calcium channel blockers such as nifedipine and verapamil, and the antihistamine cyproheptadine. Interestingly, cyproheptadine is currently being investigated in COVID-19 (NCT04820751) due to its function as an anti-serotonergic, with the possibility of improving organ dysfunction due to elevated plasma serotonin levels leading to platelet dysfunction^33–35^. Calcium channel blockers can also inhibit platelet aggregation^36^. Drug-gene interactions of these three drugs showed that they might interact with early upregulated DE genes that were involved in “Platelet degranulation” including *CLU*, *FN1*, *IGF1*, *PF4*, *PPBP*, and *TIMP1*. Thus, these drugs specific to the early phase may be targeting genes involved in platelet activation, which was detected to be elevated early in disease (**Figure 1**). As patients progressed to the “Severe” phase, the antifungals miconazole and ketoconazole, which also have immunomodulatory functions^37^, were uniquely strongly enriched. At the “Critical” phase, metformin, an anti-diabetic medication that influences energy metabolism, was the top hit, and observational studies have linked metformin use to decreased COVID-19 severity^38^. Only one drug was uniquely enriched at the Recovery phase, the anti-epileptic lacosamide.

**Figure 2.**
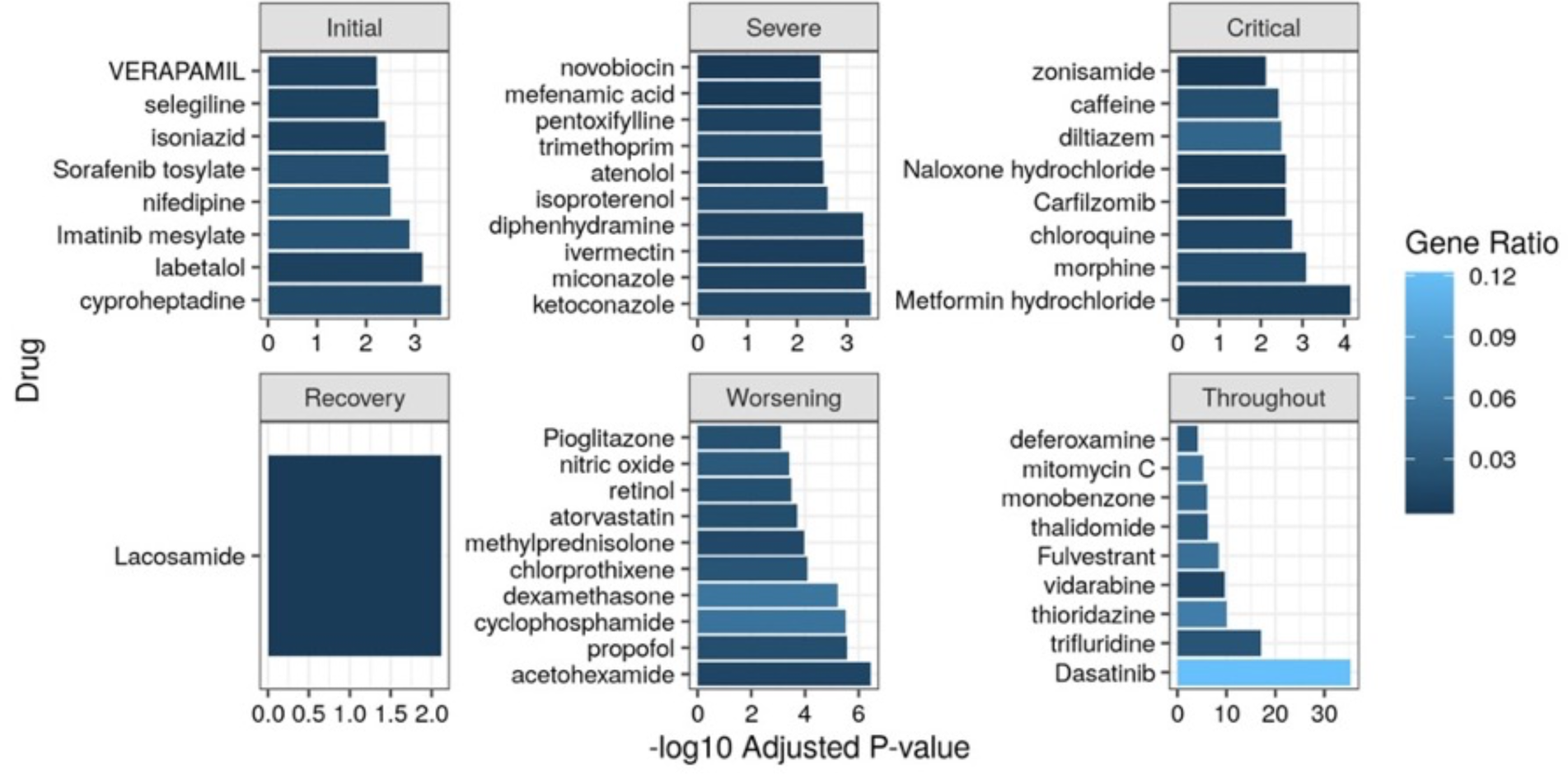
Significantly enriched drug-gene interactions from DSigDB. Bar plots indicate -log_10_ adjusted P-value and shading indicates the gene ratio. Up to 10 of the top drugs based on smallest P-value are plotted. Drugs that were uniquely enriched in the Initial stage (Mild and Moderate phases), Severe, Critical, and Recovery phase are shown (Discharge phase had no uniquely enriched drugs), as well as drugs enriched in the four “Worsening” phases (Mild, Moderate, Severe, Critical) or all six phases (“Throughout”). Gene ratio describes the proportion of DE genes found in the drug-gene interaction set.

Drugs with signatures that were enriched at multiple phases could potentially be useful administered at multiple points in the disease timeline. Notably, in the “Worsening” category (*i.e.*, drugs targeted to DE genes enriched from the Mild to Critical phases), anti-diabetic medications such as acetohexamide and pioglitazone were enriched. These anti-diabetic medications and metformin might target the metabolic dysfunction observed during increased severity of disease (**Figure 1A)**. Dexamethasone was also enriched in this category, and this medication is known to be effective in hospitalized COVID-19 patients^39^. Dasatinib, a tyrosine kinase inhibitor for chronic myeloid leukemia, was the most highly enriched drug that was enriched at all phases (“Throughout”). Interestingly, dasatinib was able to eliminate virus-induced senescent cells in COVID-19 to reduce inflammation and mitigate lung disease in animal models^40^.

### Gene signatures classified patients into disease stages for prognostication

The appropriate use of future time-dependent precision therapies is predicated on the assumption that clinicians can assess whether patients are worsening and require further support, or conversely on a recovery trajectory. Relying purely on SOFA scores is insufficient since, for example, Recovery phase samples overlap with both Moderate and Severe phase samples in terms of their SOFA score ranges. Furthermore, when patients first present to the emergency department or are admitted to the ward or ICU, there are no previous measurements to determine whether a patient is on a worsening or recovering trajectory. Thus, determining where a patient is on the disease timeline requires identifying the processes and genes uniquely dysregulated at different disease stages. The six phases were grouped into three stages for sufficient samples at each stage (Initial, Peak, and Convalescence). Indeed, the PCA separation was more pronounced when analyzing based on disease stage namely Initial (Mild and Moderate), Peak (Severe and Critical), and Convalescence (Recovery and Discharge) (**Figure S4**). Comparing each stage to the two other stages identified DE genes unique to that stage, enabling development of gene signatures that could classify patients into disease stages.

There was a progression in stage specific DE genes ranging from Initial (911) to Peak (2,014) to Convalescence (1,398), reflecting pathways (**Figure 3A**) similar to that observed during the different phases (**Figure 1**). Based on these substantial differences between stages, specific DE gene-expression signatures were developed to classify patient samples into one of the three stages. A preliminary gene signature was developed by using the top 50 upregulated DE genes for each stage relative to all other stages (**Table S3**). GSVA was performed using this preliminary gene signature to then classify patients, which it was able to do so with reasonable accuracies of 75.3%, 80.3%, and 77.7% for the Initial, Peak, and Convalescence stages, respectively (**Figure 3B**). The condensed signature with 19, 10, and 20 genes (**Table 2**) performed slightly better with accuracies of 76.7%, 81.0%, and 77.7% for Initial, Peak, and Convalescence stages, respectively (**Figure 3C, D**). Notably, these signatures were not simply detecting severity since patients in the Initial and Convalescence stages had similar SOFA scores (**Figure S2**), but also reflected the temporal aspects and trajectory of disease (**Figure S1**).

**Figure 3.**
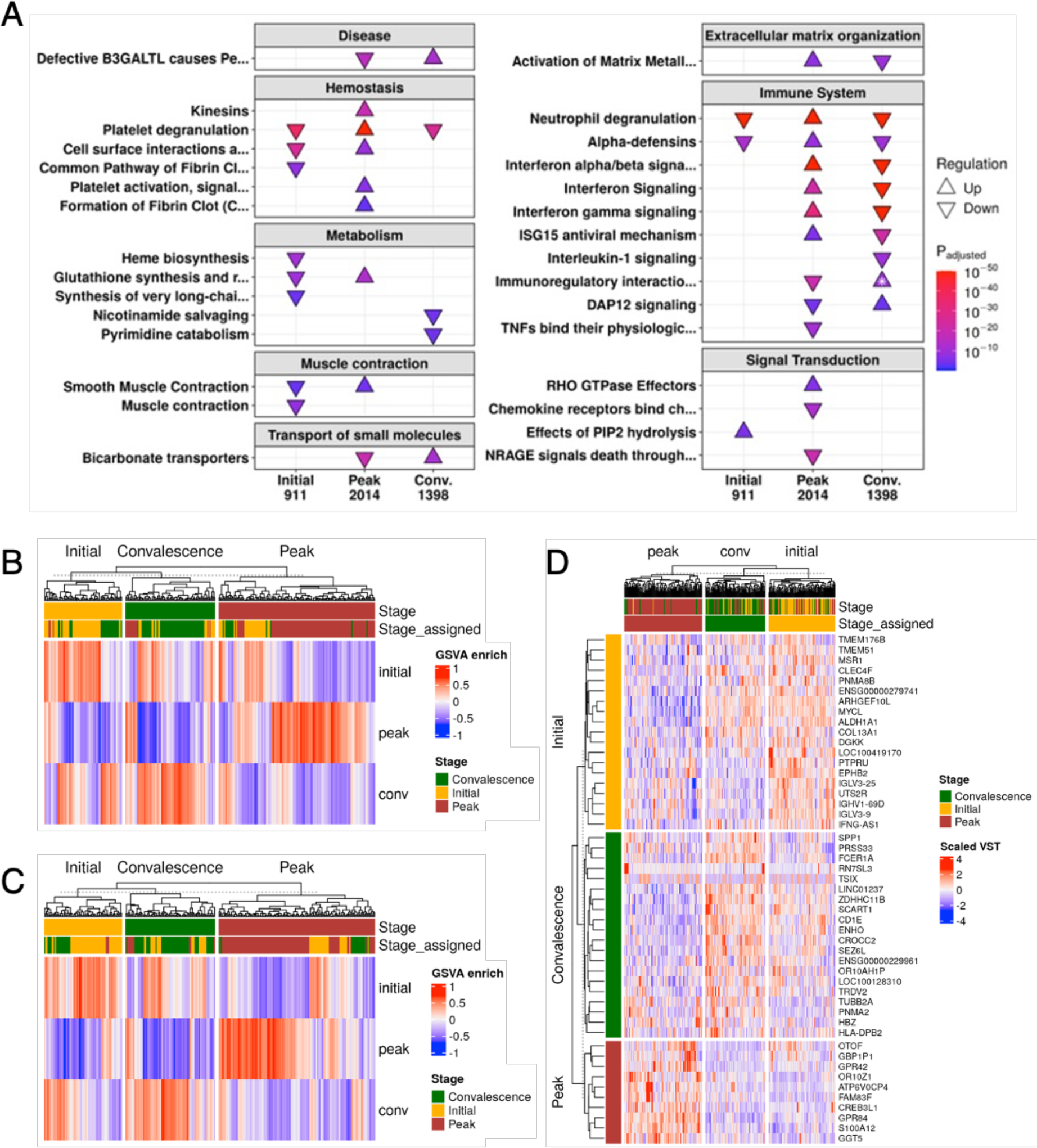
Each disease stage had different underlying mechanisms and gene signatures relative to other stages. **A:** Enriched Reactome pathways from DE genes between samples from each stage compared to samples not in that stage are shown. For one pathway, both directions were enriched (indicated by *); the direction with the lower adjusted p-value is shown. The total number of DE genes in each comparison is shown under the label. Conv: Convalescence. **B:** GSVA enrichment scores using preliminary gene signatures of Initial, Peak, and Convalescence stages for each sample. The original stage and the stage assigned using GSVA are labelled on the top bars. **C:** GSVA enrichment scores using condensed gene signatures of Initial, Peak, and Convalescence stages for each sample. **D:** Scaled variance-stabilized transformed counts of each gene in the condensed signatures.

**Table 2.** Genes and fold changes specific to each stage’s condensed gene signature. Fold changes in bold indicate genes significantly upregulated in that stage relative to all other stages, while fold changes in italics indicate non-significant fold changes from DESeq2. The full preliminary signature is found in **Table S3**. PG = pseudogene, Conv = Convalescence.

| Gene | Fold Change |  |  | Description |
| --- | --- | --- | --- | --- |
|  | Initial | Peak | Conv |  |
| *IGLV3-25* | **2.55** | *-1.49* | *-1.91* | immunoglobulin lambda variable 3-25 |
| *CLEC4F* | **2.10** | *-3.92* | *1.78* | C-type lectin domain family 4 member F |
| *COL13A1* | **1.80** | *-2.69* | *1.73* | collagen type XIII alpha 1 chain |
| *IGHV1-69D* | **1.72** | *-1.41* | *-1.39* | immunoglobulin heavy variable 1-69D |
| *TMEM176B* | **1.67** | *-1.32* | *-1.23* | transmembrane protein 176B |
| *PNMA8B* | **1.65** | *-1.89* | *1.29* | PNMA family member 8B |
| *MSR1* | **1.65** | *-1.23* | *-1.30* | macrophage scavenger receptor 1 |
| *ALDH1A1* | **1.62** | *-1.87* | *1.28* | aldehyde dehydrogenase 1 family member A1 |
| *LOC100419170* | **1.62** | *-1.26* | *-1.23* | toll like receptor 2 PG |
| *PTPRU* | **1.60** | *-1.16* | *-1.39* | protein tyrosine phosphatase receptor type U |
| *ARHGEF10L* | **1.59** | *-2.01* | *1.41* | Rho guanine nucleotide exchange factor 10 like |
| *IGLV3-9* | **1.58** | *-1.02* | *-1.64* | immunoglobulin lambda variable 3-9 |
| *DGKK* | **1.57** | *-1.85* | *1.32* | diacylglycerol kinase kappa |
| *ENSG00000279741* | **1.57** | *-1.78* | *1.25* | uncategorized |
| *IFNG-AS1* | **1.57** | *-1.65* | *1.16* | IFNG antisense RNA 1 |
| *MYCL* | **1.54** | *-1.91* | *1.38* | MYCL proto-oncogene, bHLH transcription factor |
| *UTS2R* | **1.53** | *-1.28* | *-1.13* | urotensin 2 receptor |
| *EPHB2* | **1.51** | *-1.06* | *-1.44* | EPH receptor B2 |
| *TMEM51* | **1.51** | *-1.23* | *-1.18* | transmembrane protein 51 |
| *ATP6V0CP4* | *-10.6* | **11.0** | *-4.20* | ATPase H<sup>+</sup> transporting V0 subunit c PG 4 |
| *CREB3L1* | *-4.76* | **6.54** | *-4.44* | cAMP responsive element binding protein 3 like 1 |
| *GPR42* | *-2.35* | **5.10** | *-5.98* | G protein-coupled receptor 42 |
| *GGT5* | *-5.43* | **4.08** | *-2.57* | gamma-glutamyltransferase 5 |
| *S100A12* | *-2.75* | **3.63** | *-2.64* | S100 calcium binding protein A12 |
| *GPR84* | *-2.30* | **3.58** | *-3.16* | G protein-coupled receptor 84 |
| *OR10Z1* | *-4.35* | **3.53** | *-1.84* | olfactory receptor family 10 subfamily Z member 1 |
| *GBP1P1* | *-1.28* | **3.32** | *-7.84* | guanylate binding protein 1 PG 1 |
| *OTOF* | *-1.22* | **2.95** | *-5.17* | otoferlin |
| *FAM83F* | *-3.01* | **2.87** | *-1.66* | family with sequence similarity 83 member F |
| *TSIX* | *-3.66* | *-2.35* | **6.11** | TSIX transcript, XIST antisense RNA |
| *RN7SL3* | *-3.14* | *1.02* | **4.72** | RNA component of signal recognition particle 7SL3 |
| *CROCC2* | *1.48* | *-5.50* | **3.14** | ciliary rootlet coiled-coil, rootletin family member 2 |
| *SEZ6L* | *1.13* | *-2.38* | **2.22** | seizure related 6 homolog like |
| *OR10AH1P* | *1.34* | *-2.69* | **2.19** | olfactory receptor fam. 10 subfamily AH member 1 PG |
| *CD1E* | *1.34* | *-2.81* | **2.17** | CD1e molecule |
| *FCER1A* | *-1.01* | *-2.01* | **2.14** | Fc fragment of IgE receptor Ia |
| *PNMA2* | *-3.20* | *-1.08* | **2.13** | PNMA family member 2 |
| *HBZ* | *-1.78* | *-1.38* | **2.07** | hemoglobin subunit zeta |
| *ZDHHC11B* | *1.37* | *-2.53* | **1.97** | zinc finger DHHC-type containing 11B |
| *ENHO* | *1.49* | *-2.79* | **1.97** | energy homeostasis associated |
| *TRDV2* | *-1.25* | *-1.58* | **1.95** | T cell receptor delta variable 2 |
| *PRSS33* | *-1.49* | *-1.38* | **1.95** | serine protease 33 |
| *ENSG00000229961* | *1.11* | *-1.99* | **1.92** | novel SLAM family member PG |
| *TUBB2A* | *-3.10* | *1.03* | **1.89** | tubulin beta 2A class IIa |
| *SCART1* | *1.16* | *-2.03* | **1.88** | scavenger receptor FM expressed on T cells 1 |
| *HLA-DPB2* | *-1.40* | *-1.39* | **1.87** | major histocompatibility complex, class II, DP β2 PG |
| *SPP1* | *1.02* | *-1.72* | **1.80** | secreted phosphoprotein 1 |
| *LINC01237* | *1.11* | *-1.87* | **1.79** | long intergenic non-protein coding RNA 1237 |
| *LOC100128310* | *1.17* | *-1.95* | **1.78** | uncharacterized LOC100128310 |

The condensed signatures for the Peak and Convalescence stage (based on ICU patients) were then assessed using a publicly available dataset of 42 COVID-19 and non-COVID-19 sepsis patients with samples collected at ICU admission and approximately one week later in the ICU^5^ (**Figure S9A**). In this cohort, for non-survivors, 89% (8/9) were classified as Peak at D1 and 67% (6/9) were still classified as Peak at D7, consistent with the observation that these patients had persistent immune dysfunction that led to their eventual demise (**Figure S9B**). Conversely, for survivors, 64% (21/33) of patients were classified as Peak at D1, and this significantly decreased to 24% (8/33) by D7 (p = 0.003), consistent with the observation that these patients were on a recovery trajectory and eventually discharged (**Figure S9B**). Thus, these signatures were able to identify patients at different stages of their disease in a separate external cohort.

### Disease phases were highly associated with different sepsis endotypes

Heterogeneity in patients has been attributed to many different variables. One successful approach for addressing heterogeneity is to separate patients into endotypes based on their underlying mechanistic differences. We previously identified five endotypes in early sepsis patients in the emergency room^9^, namely Neutrophilic-Suppressive (NPS), Inflammatory (INF), Interferon (IFN), Adaptive (ADA), and Innate Host Defense (IHD). The NPS and INF endotypes are associated with the worst outcomes^9^, and these endotypes have also been validated in COVID-19 patients^41^. We then investigated whether these endotypes correlated with different disease phases to determine how patients may progress through different endotypes during their disease.

Using GSVA, the enrichment score for each endotype was calculated and endotypes were assigned to each sample based on the highest enrichment score (**Figure 4**). There was a significant association between the phase and endotype to which a sample was assigned (Chi-squared p = 4.22×10^−7^) (**Figure 4C**). Specific associations were further investigated by analyzing the Chi-squared residuals, which represent positive or negative associations between the endotypes and phases. The NPS endotype, the endotype with the poorest outcome^9^, was positively associated with the Severe and Critical phases; conversely, the low severity IHD endotype was strongly positively associated with the Discharge phase and control (Follow-up) samples (**Figure 4C**). The IFN endotype was more associated with the earlier Mild and Moderate phases, likely reflecting the early antiviral response. These patterns were further reflected in the enrichment score trends (**Figure 4D**). For example, the low severity IHD endotype was significantly decreased at all phases relative to controls, with the most substantial and significant decrease at the Critical phase, while the high severity NPS endotype showed the opposite trend. Overall, the significant association between endotypes and phases was consistent with the conclusion that a portion of the heterogeneity captured by these endotypes in patients might be driven by patients presenting at different stages of their disease.

**Figure 4.**
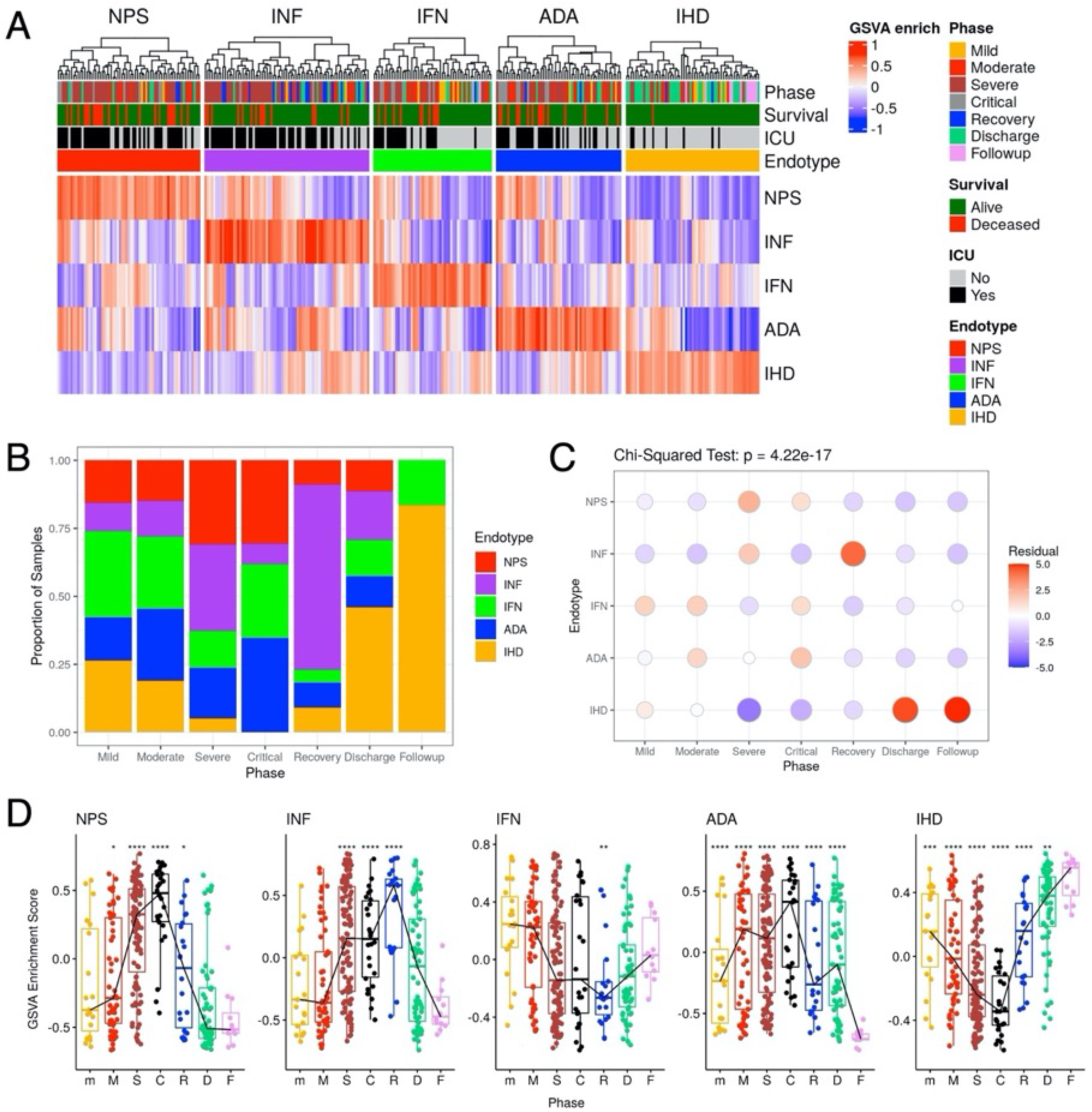
Sepsis endotypes were highly correlated with different COVID-19 disease phases. **A:** GSVA enrichment scores of each endotype signature for all samples. Samples were classified into endotypes based on the highest enrichment score. **B:** Proportion of samples with each endotype for each phase. **C:** Chi-squared residual plot of each endotype to each phase. Standardized residuals represent the association between endotypes and phases, where positive residuals indicate a positive association (red) while negative residuals indicate a negative association (blue) between a phase and an endotype. The sizes of points represent the absolute magnitude of the residual. **D:** GSVA enrichment score for each endotype signature at each phase. The trend line connects median enrichment scores for each phase. A Wilcox test was performed for each phase relative to the Followup controls. * = p<0.05, ** = p<0.01, *** = p<0.001, **** = p<0.0001. Phases were abbreviated as their first letter except for m = Mild.

## Discussion

The dynamic nature of COVID-19 can be better understood by breaking down a disease into separate phases through which individual patients can transition, which each phase reflecting mechanistic differences that require different therapeutic management. In this study, we identified six disease phases of COVID-19 based on SOFA score severity, ICU admission, and general trajectories of severity (*e.g.*, SOFA score on a downward trend or subsequent ICU discharge). Each phase was associated with distinct DE genes and pathways (**Figure 1**), as well as a variety of clinical variables (**Table S1**), strongly suggesting that the phases capture distinct mechanistic phases of illness in COVID-19. It was evident that different underlying mechanisms were initiated early or later in COVID-19, and some were ongoing even close to discharge. The heterogeneity in the host response at these different disease phases emphasized the importance of factoring time into studying dynamic diseases such as severe COVID-19. Given the known relationship of severe COVID and all-cause sepsis^7,8^, many of the findings of our study likely relate to sepsis in general.

When looking at pathway enrichment results, antiviral responses were initiated early and then were no longer evident during recovery (**Figures 1, S7**), highlighting the importance of early antiviral therapy (*e.g.*, remdesivir, monoclonal antibodies, and Paxlovid), which becomes less effective later on^42–45^. Conversely, heme metabolism was activated later, consistent with other investigations^5^, and could illustrate the importance of heme metabolism during the Recovery phase as a potential mechanism of decreasing the inflammatory response and reducing oxidative stress in myeloid cells^46^. Immune dysregulation with both heightened inflammation and adaptive suppression became very prominent later in disease and was particularly pronounced during the Critical phase of illness (**Figures 1, S7**). Immune dysregulation was even more pronounced in the Critical phase in patients who eventually died (**Figure S8**). Thus, immunomodulatory therapies are critical in targeting this immune dysregulation when it is too late for antiviral therapies to be effective.

Platelet degranulation appeared to be enriched throughout the disease timeline, peaking during the Critical phase (**Figures 1, S7**); however, this trend was the opposite of actual platelet counts, which were lowest during the Critical phase (**Table S1**) and even lower for patients who eventually died (**Figure S8, Table S2**), possibly due to excessive degranulation. The relationship between low platelets and mortality is supported by the literature, wherein thrombocytopenia in COVID-19 is associated with increased mortality in COVID-19^47^, making it a potential prognostic biomarker. Platelet activation could also be an early therapeutic target, with gene-drug signature enrichment highlighting calcium channel blockers and cyproheptadine as drugs that interact with dysregulated genes involved in platelet activation in the Mild and Moderate phases (**Figure 2**). Interestingly, hemostatic pathways were persistently enriched even through to the Discharge phase (**Figure 1**), which hints at its potential implications in “long COVID”; indeed, recent studies^48,49^ have uncovered hemostatic irregularities in patients suffering from long COVID.

Aside from possible platelet inhibitors, multiple other drugs with immunomodulatory activity were also predicted based on their enrichment in the more severe phases as well as throughout disease, including the corticosteroid dexamethasone, the tyrosine kinase inhibitor dasatinib, and antidiabetic/metabolic drugs such as metformin, acetohexamide, and pioglitazone (**Figure 2**). Dasatinib was the most enriched drug with the highest gene ratio, highlighting its effect on multiple dysregulated genes, and it would be worthwhile to further investigate its utility in severe sepsis and COVID-19. Baricitinib, another tyrosine kinase inhibitor, has shown efficacy in COVID-19^50^, suggesting the class of tyrosine kinase inhibitors could be a useful trove of potentially efficacious medications. Interestingly, the observed lower rate of COVID-19 infection in patients on tyrosine kinase inhibitors for chronic myeloid leukemia suggests potential preventative mechanisms of this drug class^51^. Dasatinib is an inhibitor of ABL kinase and Src family kinases, and both kinases are involved in viral replication of SARS-CoV-1 and SARS-CoV-2^52,53^, as well as in production of pro-inflammatory cytokines and lung fibrosis^53^. Dasatinib may confer benefits in patients by inhibiting these detrimental processes during COVID-19; indeed, dasatinib has been shown to provide survival benefits in a mouse model of sepsis^54^. Thus, this is a promising repurposed drug candidate for severe COVID-19/sepsis therapy. Antidiabetic medications should also be investigated further as COVID-19 therapies. Consistent with this, metformin and sulfonylureas (*e.g.*, acetohexamide) were shown in a meta-analysis to reduce the risk of mortality in COVID-19 patients with type 2 diabetes^55^, while pioglitazone, a thiazolidinedione, has immunomodulatory effects in addition to antidiabetic functions due to its role as a PPARɣ agonist^56^. The impact of these medications might be due to both their effects on control of glucose metabolism in combination with their immunomodulatory effects^57,58^, targeting both dysregulated metabolism and immune function during severe disease described previously (**Figure 1**). Overall, this analysis uncovered many potential repurposed drugs, with validation in the form of some top hits already currently being investigated (metformin, cyproheptadine, dasatinib) or having already shown efficacy (dexamethasone), as well as drugs that were differentially enriched in early vs. later phases. These findings are a potential starting point for identifying therapies that could work better early or later in the disease, providing personalized medicine to patients. Future clinical studies are needed to evaluate some of these therapies in COVID-19 patients. Nevertheless, further investigations and observational trials need to be performed to test these potential repurposed and affordable therapies.

Gene signatures with 10-20 genes were also identified and validated in an external cohort that could stratify patients into Initial, Peak, and Convalescence stages of disease for patient prognostication and potentially guiding new time- and disease phase-dependent personalized therapies (**Figure 3**). We also explored whether gene signatures of sepsis/COVID-19 endotypes captured different stages of disease. This was indeed found to be the case, with endotypes significantly associating with different disease phases (**Figure 4**). The poor prognosis NPS endotype was largely associated with the Severe and Critical phases, while the low severity IHD endotype was largely associated with the Discharge phase and controls. The IFN (Interferon) endotype was most associated with the Mild and Moderate phases, likely reflecting the early antiviral response.

Overall, by using a different longitudinal approach, temporal changes in key processes were identified, including early antiviral response, late heme metabolism, and severe immune, platelet, and metabolic dysregulation in high severity patients. In addition, a variety of potential disease-phase specific COVID-19 therapeutics were identified, including calcium channel blockers, cyproheptadine, dasatinib, and anti-diabetic medications. These need further clinical investigation to assess their efficacy. To facilitate time-dependent therapy, gene signatures were identified that could accurately classify patients into specific disease stages. Lastly, sepsis/COVID-19 endotypes were highly associated with specific phases, suggesting that endotypes are also capturing disease stage as part of disease heterogeneity. By understanding how patients change in their gene expression profiles over time, there is a valuable opportunity to risk stratify patients more accurately and provide time-dependent personalized treatments to intervene before patients transition to a worse phase, or to accelerate patients to recovery.

## Data Availability Statement

The datasets generated for this study can be found on GEO: GSE221234 and GSE222253. Code is available upon request (Dr. Robert Hancock).

## Ethics Statement

The Clinical Research Ethics Board of the University of British Columbia (UBC; approval number H17-01208) and Comité d’éthique de la recherche du Centre hospitalier de l’Université de Montréal (CHUM; approval numbers MP-02-2020-8929 and 19.389) provided ethics approval for all sequencing and bioinformatics studies, carried out in a manner blinded to patient identity, and written informed consent was obtained from all participants or, when incapacitated, their legal guardian before enrollment and sample collection.

## Author Contributions

RH and RL conceived the study. AA, AB, and RH contributed to the study design. AA performed bioinformatics analysis and wrote the initial draft of the paper. AA, AB, PZ, TB, JG, AL, and RH contributed to interpretation of data. DK and RL coordinated and were directly involved in sample and patient metadata collection in hospitals. AA, AB, TB, EA, PZ, and AL verified the quality and accuracy of sequencing and clinical data. RH, AL, and RL were responsible for obtaining funding. RH led the study and extensively edited the manuscript. All authors have read, edited, and approved the final version of the manuscript.

## Acknowledgments

The authors gratefully acknowledge support from the Canadian Institutes for Health Research (CIHR), the Biobanque Québécoise COVID-19, the Fonds de recherche du Québec - Santé, Génome Québec and the Public Health Agency of Canada. The authors deeply thank all the patients and their families who made this research possible.

