## Supplemental Data for "Dynamic gene expression analysis reveals distinct severity phases of immune and cellular dysregulation in COVID-19"

**Supplemental Tables**

Table S1. Clinical metadata of samples from each of the six phases.

The Kruskal-Wallis test was used, and the mean ± standard deviation of the variable is displayed, with the number of patients assessed in brackets. Significant P-values (p <0.05), indicating there were significant differences between the different phases, are bolded.

| **Clinical Metadata** | **Mild (19)** | **Moderate**  **(53)** | **Severe**  **(119)** | **Critical**  **(26)** | **Recovery**  **(22)** | **Discharge**  **(61)** | **P-value** |
| --- | --- | --- | --- | --- | --- | --- | --- |
| Temperature  (Celsius) | 36.6 ± 0.7 (19) | 36.7 ± 0.5 (53) | 37.1 ± 0.7 (119) | 36.9 ± 0.6 (26) | 37.1 ± 0.7 (22) | 36.5 ± 0.5 (60) | **<0.001** |
| Systolic BP  (mmHg) | 139.6 ± 17.8 (19) | 128.1 ± 19.4 (53) | 130.4 ± 23 (115) | 129.4 ± 22.2 (24) | 127.5 ± 15.3 (22) | 124.9 ± 19.2 (60) | 0.104 |
| Diastolic BP  (mmHg) | 79.5 ± 8.5 (19) | 73.8 ± 12.5 (53) | 59.5 ± 12.1 (115) | 54.6 ± 8.9 (24) | 63 ± 15.2 (22) | 74.3 ± 9.9 (60) | **<0.001** |
| Respiratory Rate (Breaths/min) | 19.9 ± 0.5 (19) | 20.2 ± 2  (53) | 24.8 ± 6.4 (115) | 24.8 ± 7.4 (24) | 23.5 ± 5.1 (22) | 19.5 ± 1.2 (59) | **<0.001** |
| Heart Rate  (Beats/min) | 84.2 ± 13.7 (19) | 79.2 ± 16.7 (53) | 81.5 ± 20.4 (115) | 79.1 ± 10.4 (24) | 86 ± 20.4 (22) | 76.3 ± 12.3 (60) | 0.212 |
| Leukocytes  (10^3^ cells/µL) | 8.4 ± 3.4  (19) | 7.8 ± 3.6  (46) | 9.6 ± 5  (112) | 10.7 ± 5.7 (25) | 9.5 ± 4.3  (21) | 7.3 ± 2.6  (45) | **0.031** |
| Neutrophils  (10^3^ cells/µL) | 5.8 ± 2.7  (19) | 5.7 ± 3.3  (45) | 7.7 ± 4.4 (112) | 8.6 ± 4.7  (25) | 6.6 ± 3.9  (20) | 4.6 ± 2.1  (44) | **<0.001** |
| Lymphocytes  (10^3^ cells/µL) | 1.5 ± 0.9  (19) | 1.4 ± 0.7  (45) | 0.9 ± 0.5 (112) | 0.8 ± 0.7  (25) | 1.5 ± 0.6  (20) | 1.6 ± 0.7  (44) | **<0.001** |
| Monocytes  (10^3^ cells/µL) | 0.9 ± 0.5  (19) | 0.7 ± 0.3  (45) | 0.5 ± 0.4 (112) | 0.4 ± 0.3  (25) | 0.6 ± 0.3  (20) | 0.7 ± 0.3  (44) | **<0.001** |
| Eosinophils  (10^3^ cells/µL) | 0.2 ± 0.2  (18) | 0.1 ± 0.1  (43) | 0.1 ± 0.2  (92) | 0.1 ± 0.2  (15) | 0.4 ± 0.5  (18) | 0.1 ± 0.2  (36) | **0.021** |
| Platelets  (10^3^/µL) | 312 ± 95  (19) | 282 ± 129 (46) | 297 ± 131 (112) | 180 ± 135 (25) | 348 ± 176 (21) | 317 ± 161 (45) | **<0.001** |
| Hemoglobin  (g/L) | 124 ± 16  (19) | 123 ± 19  (46) | 101 ± 19 (112) | 92 ± 18  (25) | 96 ± 16  (21) | 122 ± 23  (45) | **<0.001** |
| Urea  (mg/dL) | 7 ± 2.2  (12) | 7.5 ± 4  (25) | 11.1 ± 7.6 (112) | 15 ± 11  (25) | 8.6 ± 5.6  (21) | 6.4 ± 3.4  (27) | **<0.001** |
| Creatinine  (µmol/L) | 71.9 ± 15.6 (18) | 86.5 ± 60.9 (46) | 129.3 ± 215 (112) | 394 ± 293 (25) | 88.2 ± 84.1 (21) | 71.2 ± 27.8 (46) | **<0.001** |
| Glucose  (mmol/L) | 6.4 ± 2.3  (14) | 7.4 ± 3.3  (43) | 7.6 ± 2.2 (116) | 8.2 ± 2.9  (25) | 8 ± 2.3  (20) | 7.5 ± 3.3  (38) | 0.072 |
| Sodium  (mmol/L) | 138.7 ± 4.6 (18) | 140.6 ± 4 (46) | 140.9 ± 5.1 (114) | 137.5 ± 3.8 (25) | 141 ± 3.4 (21) | 135.6 ± 18.3 (44) | **<0.001** |
| Potassium  (mmol/L) | 4 ± 0.3  (18) | 4 ± 0.4  (46) | 4.1 ± 2.7 (113) | 4.1 ± 0.4  (25) | 4 ± 0.4  (21) | 4.1 ± 0.4  (43) | 0.120 |
| AST (units/L) | 49.5 ± 32  (6) | 29.2 ± 12.1 (10) | 48.7 ± 33.6 (99) | 163 ± 416 (24) | 116 ± 328 (20) | 35.1 ± 18.1 (18) | **0.032** |
| ALT  (units/L) | 57 ± 59.8 (11) | 41.5 ± 27.6 (16) | 54.2 ± 54.5 (100) | 76.6 ± 168 (25) | 132 ± 362 (20) | 75.8 ± 73.5 (24) | 0.264 |
| Total Bilirubin  (µmol/L) | 10.9 ± 3  (8) | 11.5 ± 8.4 (11) | 11.9 ± 9 (100) | 30.8 ± 37.9 (25) | 10.2 ± 4.7 (20) | 9.2 ± 3.4  (20) | 0.278 |
| C-reactive Protein  (mg/L) | 72.2 ± 69.4 (12) | 57.8 ± 49.2 (40) | 165 ± 105 (53) | 221 ± 121  (9) | 34.6 ± 29.7 (6) | 38.4 ± 35.3 (25) | **<0.001** |
| Procalcitonin  (ng/mL) | 0.1  (1) | 0.3 ± 0.3  (14) | 0.7 ± 0.6  (7) | 20.7 ± 16.3 (3) | NA  (0) | 0.1 ± 0.1  (7) | **0.006** |
| Venous Lactate  (mmol/L) | 1.5  (1) | 0.8 ± 0.2  (2) | 1.4 ± 0.7 (107) | 1.2 ± 0.7  (25) | 1.2 ± 0.3  (19) | 1.2 ± 0.3  (3) | 0.416 |
| D-dimer  (ng/mL) | 681  (1) | 2272 ± 797 (4) | 2479 ± 2306 (28) | 2284 ± 2823 (7) | 2180 ± 1800 (4) | 1075  (1) | 0.552 |
| Fibrinogen  (g/L) | NA  (0) | 5.7 ± 0.6  (3) | 6.3 ± 2.5  (53) | 6.7 ± 1.4  (17) | 5.4 ± 1.2  (10) | NA  (0) | 0.245 |
| Ferritin  (ng/mL) | 564  (1) | 518 ± 527  (3) | 853 ± 1196 (11) | NA  (0) | 1595 ± 2658 (3) | 685 ± 470  (6) | 0.912 |
| Total SOFA Score | 1 ± 0  (19) | 2.8 ± 1.2  (53) | 7.9 ± 2.4 (119) | 13 ± 0.9  (26) | 6 ± 2.6  (22) | 1.4 ± 0.8  (61) | **<0.001** |
| Respiratory | 1 ± 0  (19) | 2.1 ± 0.4  (53) | 2.9 ± 0.8 (119) | 3 ± 0.7  (26) | 2.3 ± 0.7  (22) | 1.2 ± 0.5  (61) | **<0.001** |
| Renal | 0 ± 0  (19) | 0.2 ± 0.6  (53) | 0.6 ± 1.1 (119) | 2.6 ± 1.3  (26) | 0.5 ± 1.1  (22) | 0 ± 0.3  (61) | **<0.001** |
| Cardiovascular | 0 ± 0  (19) | 0.1 ± 0.3  (53) | 1.3 ± 1  (119) | 2 ± 0  (26) | 1 ± 1  (22) | 0 ± 0  (61) | **<0.001** |
| Neurological | 0 ± 0  (19) | 0.1 ± 0.3  (53) | 2.8 ± 1.6 (119) | 3.9 ± 0.3  (26) | 2.2 ± 1.7  (22) | 0 ± 0.3  (61) | **<0.001** |
| Hepatic | 0 ± 0  (19) | 0 ± 0.3  (53) | 0.2 ± 0.5 (119) | 0.7 ± 1  (26) | 0 ± 0  (22) | 0 ± 0  (61) | **<0.001** |
| Coagulation | 0 ± 0  (19) | 0.2 ± 0.4  (53) | 0.2 ± 0.5 (119) | 0.8 ± 0.8  (26) | 0 ± 0.2  (22) | 0 ± 0.2  (61) | **<0.001** |
| WHO Severity Scale | 4.7 ± 0.5  (19) | 5.1 ± 0.5  (53) | 7.9 ± 1.1 (119) | 8.7 ± 0.5  (26) | 7 ± 1.2  (22) | 4.8 ± 0.5  (61) | **<0.001** |

Table S2. Clinical metadata of Critical phase samples from patients who were eventually deceased or survived.

The Wilcoxon Rank-Sum test was used, and the mean ± standard deviation of the variable is displayed, with the number of patients assessed in brackets. Significant p-values (p <0.05) are bolded.

| **Clinical Metadata** | **Deceased (14)** | **Survived (12)** | **P-value** |
| --- | --- | --- | --- |
| Temperature (Celsius) | 37 ± 0.5 (14) | 36.9 ± 0.7 (12) | 0.772 |
| Systolic BP (mmHg) | 129 ± 21.9 (12) | 129.8 ± 23.5 (12) | 0.840 |
| Diastolic BP (mmHg) | 55.9 ± 9.7 (12) | 53.3 ± 8.1 (12) | 0.466 |
| Respiratory Rate (breaths/min) | 23.5 ± 8.1 (12) | 26 ± 6.7 (12) | 0.541 |
| Heart Rate (beats/min) | 79.8 ± 11.8 (12) | 78.4 ± 9.2 (12) | 0.862 |
| Leukocytes (10^3^ cells/µL) | 12.1 ± 5.2 (14) | 9 ± 6 (11) | 0.095 |
| Neutrophils (10^3^ cells/µL) | 9.6 ± 3.9 (14) | 7.2 ± 5.5 (11) | 0.218 |
| Lymphocytes (10^3^ cells/µL) | 0.8 ± 0.6 (14) | 0.8 ± 0.8 (11) | 0.956 |
| Monocytes (10^3^ cells/µL) | 0.5 ± 0.3 (14) | 0.4 ± 0.3 (11) | 0.661 |
| Eosinophils (10^3^ cells/µL) | 0.1 ± 0.2 (8) | 0.2 ± 0.1 (7) | 0.104 |
| Basophils (10^3^ cells/µL) | 0 ± 0 (14) | 0 ± 0 (11) | 0.190 |
| Platelets (10^3^ cells/µL) | 165.1 ± 165.2 (14) | 199.7 ± 86.2 (11) | **0.031** |
| Hemoglobin (g/L) | 95.2 ± 20.7 (14) | 87.7 ± 14.4 (11) | 0.493 |
| Urea (mg/dL) | 15 ± 12.8 (14) | 15 ± 8.9 (11) | 0.603 |
| Creatinine (µmol/L) | 312.9 ± 237.3 (14) | 497.1 ± 333.5 (11) | 0.090 |
| Glucose (mmol/L) | 7.3 ± 2.8 (14) | 9.2 ± 2.8 (11) | 0.100 |
| Sodium (mmol/L) | 136.9 ± 3.6 (14) | 138.2 ± 4.1 (11) | 0.602 |
| Potassium (mmol/L) | 4.2 ± 0.4 (14) | 4 ± 0.5 (11) | 0.205 |
| AST (units/L) | 244.2 ± 561.1 (13) | 67.9 ± 47.3 (11) | 0.664 |
| ALT (units/L) | 113.3 ± 220.1 (14) | 30 ± 24.7 (11) | 0.476 |
| Total Bilirubin (µmol/L) | 29.3 ± 25.9 (14) | 32.7 ± 50.7 (11) | 0.366 |
| C-reactive Protein (mg/L) | 233.8 ± 117.3 (6) | 195.7 ± 151.1 (3) | 0.697 |
| Venous lactate (mmol/L) | 1.4 ± 0.9 (14) | 1 ± 0.5 (11) | 0.146 |
| D-dimer (ng/mL) | 2540.8 ± 3414.7 (5) | 1642 ± 141.4 (2) | 0.561 |
| Fibrinogen (g/L) | 6.5 ± 1.3 (10) | 7 ± 1.5 (7) | 0.071 |
| Sampling Time (Days Post-Admission) | 7.1 ± 4.3 (14) | 5.6 ± 3.7 (12) | 0.323 |
| Total SOFA Score | 13.2 ± 1.1 (14) | 12.8 ± 0.6 (12) | 0.243 |
| Respiratory | 3.1 ± 0.7 (14) | 3 ± 0.7 (12) | 0.823 |
| Renal | 2.1 ± 1.3 (14) | 3.1 ± 1.2 (12) | 0.066 |
| Cardiovascular | 2 ± 0 (14) | 2 ± 0 (12) | 1.000 |
| Neurological | 3.9 ± 0.3 (14) | 3.9 ± 0.3 (12) | 0.956 |
| Hepatic | 0.9 ± 0.9 (14) | 0.5 ± 1 (12) | 0.273 |
| Coagulation | 1.2 ± 0.8 (14) | 0.2 ± 0.5 (12) | **0.003** |
| WHO Severity Scale | 8.6 ± 0.5 (14) | 8.7 ± 0.5 (12) | 0.925 |

**Table S3. Genes and fold changes specific to each cluster.** Fold changes in bold indicate the top 50 (top 47 for Initial) genes significantly upregulated in that stage relative to all other stages (full gene signature), while fold changes in italics indicate non-significant fold changes with an adjusted p-value >0.05 from DESeq2. Genes in bold indicate condensed signature genes from LASSO regression (**Table 2**). Conv: Convalescence.

| **Gene** | **Fold Change** | | | **Description** |
| --- | --- | --- | --- | --- |
|  | **Initial** | **Peak** | **Conv** |  |
| ENSG00000227242 | **19.8** | *-8.22* | *-3.27* | phosphodiesterase 4D interacting protein (PDE4DIP) pseudogene |
| IGHV1-2 | **3.01** | *-1.21* | -1.56 | immunoglobulin heavy variable 1-2 |
| **IGLV3-25** | **2.55** | *-1.49* | -1.91 | immunoglobulin lambda variable 3-25 |
| IGLV3-10 | **2.38** | *-1.28* | -2.14 | immunoglobulin lambda variable 3-10 |
| IGHV5-10-1 | **2.31** | *-1.36* | *-1.79* | immunoglobulin heavy variable 5-10-1 |
| **CLEC4F** | **2.10** | -3.92 | 1.78 | C-type lectin domain family 4 member F |
| IGHGP | **2.03** | *-1.14* | -1.95 | immunoglobulin heavy constant gamma P (non-functional) |
| KCTD14 | **1.96** | *1.22* | -4.26 | potassium channel tetramerization domain containing 14 |
| ENSG00000254829 | **1.91** | *1.22* | -3.94 | lncRNA |
| IGHV4-34 | **1.85** | *1.13* | -2.27 | immunoglobulin heavy variable 4-34 |
| SCGB3A1 | **1.83** | -1.71 | *1.04* | secretoglobin family 3A member 1 |
| **COL13A1** | **1.80** | -2.69 | 1.73 | collagen type XIII alpha 1 chain |
| IGHV1-24 | **1.79** | *-1.37* | *-1.23* | immunoglobulin heavy variable 1-24 |
| IGHV2-5 | **1.78** | *-1.25* | *-1.42* | immunoglobulin heavy variable 2-5 |
| IGLV2-18 | **1.75** | *-1.14* | -1.55 | immunoglobulin lambda variable 2-18 |
| IGLV3-19 | **1.72** | *1.23* | -2.31 | immunoglobulin lambda variable 3-19 |
| **IGHV1-69D** | **1.72** | *-1.41* | *-1.39* | immunoglobulin heavy variable 1-69D |
| IGHV4-31 | **1.69** | *-1.28* | *-1.27* | immunoglobulin heavy variable 4-30-2 |
| IGHG3 | **1.68** | *-1.04* | -1.74 | immunoglobulin heavy constant gamma 3 |
| **TMEM176B** | **1.67** | *-1.32* | *-1.23* | transmembrane protein 176B |
| **PNMA8B** | **1.65** | -1.89 | *1.29* | PNMA family member 8B |
| **MSR1** | **1.65** | *-1.23* | *-1.30* | macrophage scavenger receptor 1 |
| BOK | **1.62** | -1.61 | *1.10* | BCL2 family apoptosis regulator BOK |
| **ALDH1A1** | **1.62** | -1.87 | *1.28* | aldehyde dehydrogenase 1 family member A1 |
| **LOC100419170** | **1.62** | *-1.26* | *-1.23* | toll like receptor 2 pseudogene |
| AXL | **1.61** | *-1.34* | *-1.16* | AXL receptor tyrosine kinase |
| LINC01484 | **1.61** | *-1.06* | -1.56 | long intergenic non-protein coding RNA 1484 |
| **PTPRU** | **1.60** | *-1.16* | *-1.39* | protein tyrosine phosphatase receptor type U |
| NA | **1.60** | -1.77 | *1.23* | lncRNA |
| ADGRD1 | **1.60** | -2.41 | 1.66 | adhesion G protein-coupled receptor D1 |
| MYCL-AS1 | **1.60** | -2.06 | *1.42* | MYCL antisense RNA 1 |
| **ARHGEF10L** | **1.59** | -2.01 | *1.41* | Rho guanine nucleotide exchange factor 10 like |
| **IGLV3-9** | **1.58** | *-1.02* | -1.64 | immunoglobulin lambda variable 3-9 |
| **DGKK** | **1.57** | -1.85 | *1.32* | diacylglycerol kinase kappa |
| **ENSG00000279741** | **1.57** | -1.78 | *1.25* | NA |
| **IFNG-AS1** | **1.57** | -1.65 | *1.16* | IFNG antisense RNA 1 |
| DAGLA | **1.56** | -2.01 | *1.43* | diacylglycerol lipase alpha |
| FLT4 | **1.55** | -2.10 | 1.49 | fms related receptor tyrosine kinase 4 |
| IGHV3-66 | **1.55** | *-1.06* | *-1.45* | immunoglobulin heavy variable 3-66 |
| **MYCL** | **1.54** | -1.91 | *1.38* | MYCL proto-oncogene, bHLH transcription factor |
| MIR8071-1 | **1.54** | *-1.12* | *-1.36* | microRNA 8071-1 |
| PID1 | **1.53** | -2.95 | 2.06 | phosphotyrosine interaction domain containing 1 |
| **UTS2R** | **1.53** | *-1.28* | *-1.13* | urotensin 2 receptor |
| IGKV3-15 | **1.52** | *-1.16* | *-1.26* | immunoglobulin kappa variable 3-15 |
| IGLC6 | **1.52** | *1.31* | -1.68 | immunoglobulin lambda constant 6 |
| **EPHB2** | **1.51** | *-1.06* | *-1.44* | EPH receptor B2 |
| **TMEM51** | **1.51** | *-1.23* | *-1.18* | transmembrane protein 51 |
| MTCO1P40 | -4.44 | **11.6** | -2.58 | MT-CO1 pseudogene 40 |
| **ATP6V0CP4** | -10.63 | **11.0** | -4.20 | ATPase H+ transporting V0 subunit c pseudogene 4 |
| CD177 | -4.00 | **6.68** | -5.24 | CD177 molecule |
| **CREB3L1** | -4.76 | **6.54** | -4.44 | cAMP responsive element binding protein 3 like 1 |
| ENSG00000243144 | -4.79 | **6.54** | -4.35 | lncRNA |
| ENSG00000272396 | -3.89 | **6.50** | -5.13 | lncRNA |
| ZDHHC19 | -2.95 | **6.32** | -7.16 | zinc finger DHHC-type palmitoyltransferase 19 |
| SLC6A19 | -7.62 | **6.28** | -2.58 | solute carrier family 6 member 19 |
| CD177P1 | -3.12 | **5.70** | -4.89 | CD177 molecule pseudogene 1 |
| **GPR42** | -2.35 | **5.10** | -5.98 | G protein-coupled receptor 42 |
| CYP19A1 | -2.48 | **5.06** | -5.70 | cytochrome P450 family 19 subfamily A member 1 |
| ART4 | -5.94 | **5.06** | -2.31 | ADP-ribosyltransferase 4 (inactive) (Dombrock blood group) |
| A4GALT | -3.34 | **4.69** | -2.87 | alpha 1,4-galactosyltransferase (P blood group) |
| RETN | -4.06 | **4.47** | -2.75 | resistin |
| NRARP | -3.53 | **4.44** | -1.89 | NOTCH regulated ankyrin repeat protein |
| NNMT | -2.58 | **4.11** | -3.71 | nicotinamide N-methyltransferase |
| **GGT5** | -5.43 | **4.08** | -2.57 | gamma-glutamyltransferase 5 |
| PCSK9 | -2.68 | **3.86** | -2.85 | proprotein convertase subtilisin/kexin type 9 |
| FFAR3 | -1.92 | **3.84** | -4.47 | free fatty acid receptor 3 |
| KLF14 | -2.22 | **3.66** | -3.36 | Kruppel like factor 14 |
| **S100A12** | -2.75 | **3.63** | -2.64 | S100 calcium binding protein A12 |
| **GPR84** | -2.30 | **3.58** | -3.16 | G protein-coupled receptor 84 |
| **OR10Z1** | -4.35 | **3.53** | -1.84 | olfactory receptor family 10 subfamily Z member 1 |
| ARG1 | -3.18 | **3.51** | -2.19 | arginase 1 |
| CCNA1 | -1.96 | **3.48** | -3.46 | cyclin A1 |
| DEFA3 | -2.33 | **3.46** | -1.92 | defensin alpha 3 |
| TNFAIP8L3 | -1.95 | **3.43** | -3.53 | TNF alpha induced protein 8 like 3 |
| PRTN3 | -2.60 | **3.41** | -2.60 | proteinase 3 |
| HP | -2.75 | **3.36** | -2.41 | haptoglobin |
| WFDC1 | -2.95 | **3.36** | -2.22 | WAP four-disulfide core domain 1 |
| RHAG | -5.28 | **3.32** | -1.58 | Rh associated glycoprotein |
| **GBP1P1** | *-1.28* | **3.32** | -7.84 | guanylate binding protein 1 pseudogene 1 |
| OLFM4 | -3.10 | **3.25** | -1.95 | olfactomedin 4 |
| MMP8 | -3.20 | **3.25** | -1.97 | matrix metallopeptidase 8 |
| YPEL4 | -3.46 | **3.23** | -1.79 | yippee like 4 |
| FHDC1 | -2.53 | **3.23** | -1.89 | FH2 domain containing 1 |
| ELANE | -2.66 | **3.20** | -2.38 | elastase, neutrophil expressed |
| MCEMP1 | -2.08 | **3.16** | -2.77 | mast cell expressed membrane protein 1 |
| ERFE | -3.73 | **3.14** | -1.77 | erythroferrone |
| S100A8 | -3.56 | **3.12** | -2.33 | S100 calcium binding protein A8 |
| TMCC2 | -4.20 | **3.01** | -1.54 | transmembrane and coiled-coil domain family 2 |
| CTSG | -2.55 | **3.01** | -2.30 | cathepsin G |
| IL1R2 | -1.97 | **2.97** | -2.66 | interleukin 1 receptor type 2 |
| **OTOF** | *-1.22* | **2.95** | -5.17 | otoferlin |
| LCN2 | -2.71 | **2.93** | -1.95 | lipocalin 2 |
| ANKRD22 | -1.67 | **2.91** | -3.16 | ankyrin repeat domain 22 |
| PGLYRP1 | -2.41 | **2.89** | -2.10 | peptidoglycan recognition protein 1 |
| **FAM83F** | -3.01 | **2.87** | -1.66 | family with sequence similarity 83 member F |
| TCN1 | -2.53 | **2.85** | -1.97 | transcobalamin 1 |
| ANXA3 | -2.23 | **2.83** | -2.19 | annexin A3 |
| XIST | -3.66 | -2.39 | **6.28** | X inactive specific transcript |
| **TSIX** | -3.66 | -2.35 | **6.11** | TSIX transcript, XIST antisense RNA |
| **RN7SL3** | -3.14 | *1.02* | **4.72** | RNA component of signal recognition particle 7SL3 |
| **CROCC2** | *1.48* | -5.50 | **3.14** | ciliary rootlet coiled-coil, rootletin family member 2 |
| UICLM | *1.49* | -5.17 | **3.03** | up-regulated in colorectal cancer liver metastasis |
| CTSE | -1.93 | -1.61 | **2.77** | cathepsin E |
| LYPD2 | *1.49* | -3.66 | **2.50** | LY6/PLAUR domain containing 2 |
| ST8SIA1 | *-1.30* | -1.77 | **2.28** | ST8 alpha-N-acetyl-neuraminide alpha-2,8-sialyltransferase 1 |
| **SEZ6L** | *1.13* | -2.38 | **2.22** | seizure related 6 homolog like |
| **OR10AH1P** | *1.34* | -2.69 | **2.19** | olfactory receptor family 10 subfamily AH member 1 pseudogene |
| ADAM23 | *1.16* | -2.45 | **2.19** | ADAM metallopeptidase domain 23 |
| **CD1E** | *1.34* | -2.81 | **2.17** | CD1e molecule |
| LINC01857 | *-1.24* | -1.75 | **2.16** | long intergenic non-protein coding RNA 1857 |
| **FCER1A** | *-1.01* | -2.01 | **2.14** | Fc fragment of IgE receptor Ia |
| **PNMA2** | -3.20 | *-1.08* | **2.13** | PNMA family member 2 |
| **HBZ** | -1.78 | *-1.38* | **2.07** | hemoglobin subunit zeta |
| PID1 | 1.53 | -2.95 | **2.06** | phosphotyrosine interaction domain containing 1 |
| TIFAB | *1.42* | -2.75 | **2.03** | TIFA inhibitor |
| **ZDHHC11B** | *1.37* | -2.53 | **1.97** | zinc finger DHHC-type containing 11B |
| **ENHO** | *1.49* | -2.79 | **1.97** | energy homeostasis associated |
| LINC01259 | *1.22* | -2.25 | **1.96** | long intergenic non-protein coding RNA 1259 |
| MRC2 | -1.58 | -1.35 | **1.96** | mannose receptor C type 2 |
| ENSG00000230729 | *1.48* | -2.73 | **1.96** | lncRNA |
| NELL2 | *1.15* | -2.11 | **1.95** | neural EGFL like 2 |
| **TRDV2** | *-1.25* | -1.58 | **1.95** | T cell receptor delta variable 2 |
| ENSG00000285476 | *1.17* | -2.14 | **1.95** | NA |
| ENSG00000261655 | *1.18* | -2.14 | **1.95** | lncRNA |
| **PRSS33** | *-1.49* | *-1.38* | **1.95** | serine protease 33 |
| **ENSG00000229961** | *1.11* | -1.99 | **1.92** | novel SLAM family member pseudogene |
| SLC4A10 | *1.19* | -2.16 | **1.91** | solute carrier family 4 member 10 |
| **TUBB2A** | -3.10 | *1.03* | **1.89** | tubulin beta 2A class IIa |
| NOG | *1.21* | -2.11 | **1.89** | noggin |
| **SCART1** | *1.16* | -2.03 | **1.88** | scavenger receptor family member expressed on T cells 1 |
| LYNX1 | *1.29* | -2.22 | **1.87** | Ly6/neurotoxin 1 |
| **HLA-DPB2** | *-1.40* | *-1.39* | **1.87** | major histocompatibility complex, class II, DP beta 2 (pseudogene) |
| DBH | *1.18* | -2.03 | **1.84** | dopamine beta-hydroxylase |
| ZNF890P | *1.14* | -1.93 | **1.83** | zinc finger protein 890, pseudogene |
| ELOVL4 | *1.13* | -1.96 | **1.83** | ELOVL fatty acid elongase 4 |
| ENSG00000226945 | *1.06* | -1.83 | **1.83** | ribosomal protein L13a (RPL13A) pseudogene |
| OLFM1 | *1.31* | -2.22 | **1.82** | olfactomedin 1 |
| GCNT4 | *1.19* | -2.00 | **1.80** | glucosaminyl (N-acetyl) transferase 4 |
| ENSG00000286342 | -2.23 | *-1.06* | **1.80** | NA |
| **SPP1** | *1.02* | -1.72 | **1.80** | secreted phosphoprotein 1 |
| SEC14L3 | -1.52 | *-1.27* | **1.80** | SEC14 like lipid binding 3 |
| **LINC01237** | *1.11* | -1.87 | **1.79** | long intergenic non-protein coding RNA 1237 |
| ENSG00000226423 | *1.20* | -1.99 | **1.79** | lncRNA |
| ENSG00000286330 | *1.06* | -1.78 | **1.78** | NA |
| **LOC100128310** | *1.17* | -1.95 | **1.78** | uncharacterized LOC100128310 |
| PNMA6A | *1.09* | -1.82 | **1.78** | PNMA family member 6A |

**Table S4. Additional demographics of patients in the BQC-19 biobank.** For categorical variables, significance was tested using the Chi-squared test with Yates’s correction, or Fisher’s Exact test if any expected value was <5, and the percentage and fraction of patients fitting the category is displayed. For continuous variables, the Wilcoxon Rank-Sum test was used, and the mean ± standard deviation of the variable is displayed, with the number of patients assessed in brackets. Significant P-values (p <0.05) are bolded.

| **Clinical Metadata** | **Non-ICU (62)** | **Admitted to ICU (66)** | **P value** |
| --- | --- | --- | --- |
| **Comorbidities** | | | |
| Asthma (Yes) | 8.1% (5/62) | 18.2% (12/66) | 0.154 |
| COPD (Yes) | 12.9% (8/62) | 12.1% (8/66) | 1.000 |
| Chronic Lung Disease (Yes) | 6.5% (4/62) | 13.6% (9/66) | 0.293 |
| Atrial Fibrillation/Flutter (Yes) | 11.3% (7/62) | 10.6% (7/66) | 1.000 |
| Hypertension (Yes) | 45.2% (28/62) | 66.7% (44/66) | **0.023** |
| Heart Failure (Yes) | 8.1% (5/62) | 3.0% (2/66) | 0.263 |
| Coronary Artery Disease (Yes) | 8.1% (5/62) | 15.2% (10/66) | 0.332 |
| Chronic Cardiac Disease (Yes) | 4.8% (3/62) | 13.6% (9/66) | 0.161 |
| Liver disease (Yes) | 8.1% (5/62) | 19.7% (13/66) | 0.102 |
| Chronic Hematologic Disease (Yes) | 4.8% (3/62) | 7.6% (5/66) | 0.719 |
| Chronic Kidney Disease (Yes) | 24.2% (15/62) | 24.2% (16/66) | 1.000 |
| On Dialysis (Yes) | 1.9% (1/52) | 23.8% (5/21) | **0.007** |
| Psychiatric Disease (Yes) | 22.6% (14/62) | 7.6% (5/66) | **0.033** |
| Obesity (Yes) | 14.5% (9/62) | 28.8% (19/66) | 0.082 |
| Diabetes (Yes) | 33.9% (21/62) | 42.4% (28/66) | 0.416 |
| AIDS (Yes) | 1.6% (1/62) | 4.5% (3/66) | 0.620 |
| Immunosuppressed (Yes) | 9.7% (6/62) | 12.1% (8/66) | 0.873 |
| Rheumatologic Disease (Yes) | 0.0% (0/62) | 10.6% (7/66) | **0.013** |
| Cancer (Yes) | 12.9% (8/62) | 7.6% (5/66) | 0.481 |
| **Arrival Values** | | | |
| Temperature (Celsius) | 37 ± 0.9 (62) | 37.5 ± 1.2 (62) | 0.081 |
| Systolic BP (mmHg) | 131.9 ± 21.9 (62) | 125.1 ± 22.4 (66) | 0.097 |
| Diastolic BP (mmHg) | 77.4 ± 10.2 (62) | 69.9 ± 14 (66) | **0.001** |
| Respiratory Rate (breaths/min) | 20.9 ± 3.3 (62) | 23.9 ± 8.1 (64) | 0.089 |
| Heart Rate (beats/min) | 86.3 ± 17.1 (62) | 91.8 ± 19.6 (66) | 0.099 |
| O_2_ Saturation at Room Air | 95.4 ± 2.5 (52) | 90.8 ± 7.9 (35) | **0.001** |
| **Symptoms at Admission** | | | |
| Asymptomatic (Yes) | 8.1% (5/62) | 0.0% (0/66) | **0.024** |
| Confusion (Yes) | 11.3% (6/53) | 20.0% (13/65) | 0.306 |
| Diarrhea (Yes) | 31.5% (17/54) | 32.8% (21/64) | 1.000 |
| Abdominal Pain (Yes) | 7.4% (4/54) | 15.9% (10/63) | 0.262 |
| Chest Pain (Yes) | 11.1% (6/54) | 31.7% (20/63) | **0.014** |
| Dyspnea (Yes) | 63.0% (34/54) | 92.2% (59/64) | **<0.001** |
| Dizziness (Yes) | 9.4% (5/53) | 7.9% (5/63) | 1.000 |
| Extremity Numbness (Yes) | 13.5% (7/52) | 8.1% (5/62) | 0.529 |
| Fatigue (Yes) | 35.2% (19/54) | 58.5% (38/65) | **0.019** |
| Fever (Yes) | 42.6% (23/54) | 84.8% (56/66) | **<0.001** |
| Hemoptysis (Yes) | 1.9% (1/54) | 19.4% (12/62) | **0.007** |
| Loss of Appetite (Yes) | 24.1% (13/54) | 18.8% (12/64) | 0.632 |
| Sore Throat (Yes) | 7.3% (4/55) | 12.7% (8/63) | 0.504 |
| Headache (Yes) | 7.3% (4/55) | 14.1% (9/64) | 0.374 |
| Myalgia (Yes) | 12.7% (7/55) | 29.7% (19/64) | **0.044** |
| Nausea/Vomiting (Yes) | 20.0% (11/55) | 25.0% (16/64) | 0.667 |
| Loss of Taste/Smell (Yes) | 7.3% (4/55) | 4.8% (3/63) | 0.704 |
| Rhinorrhea (Yes) | 0.0% (0/55) | 7.9% (5/63) | 0.060 |
| Cough (Yes) | 58.2% (32/55) | 70.8% (46/65) | 0.212 |
| Aphasia/Dysphasia (Yes) | 3.6% (2/55) | 4.8% (3/63) | 1.000 |

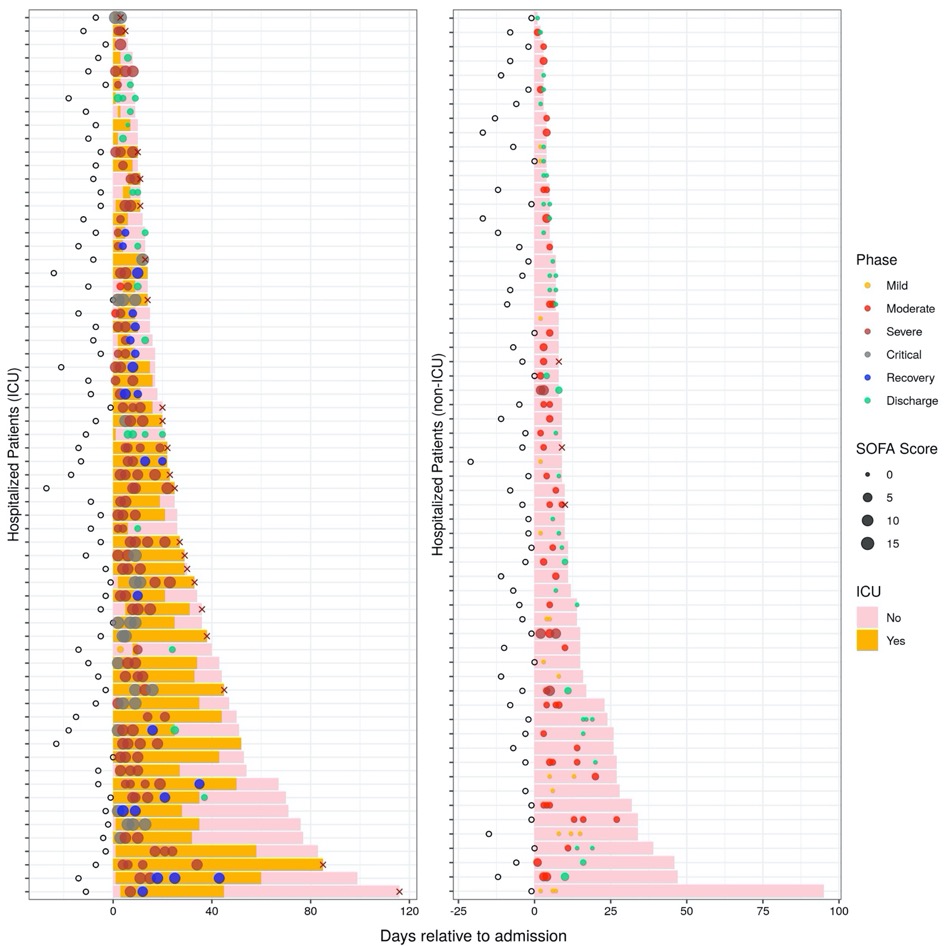

Figure S1. Disease trajectories of patients.

Bars indicate hospitalization duration of 128 patients, separated into patients admitted to the ICU at any time (left) or never (right). Circles indicate blood sampling time, with the SOFA score and assigned phase corresponding to size and colour of the point, respectively. A total of 300 samples were collected from patients. Open circles indicate when COVID-19 symptoms began, if known. X’s indicate when a patient died.

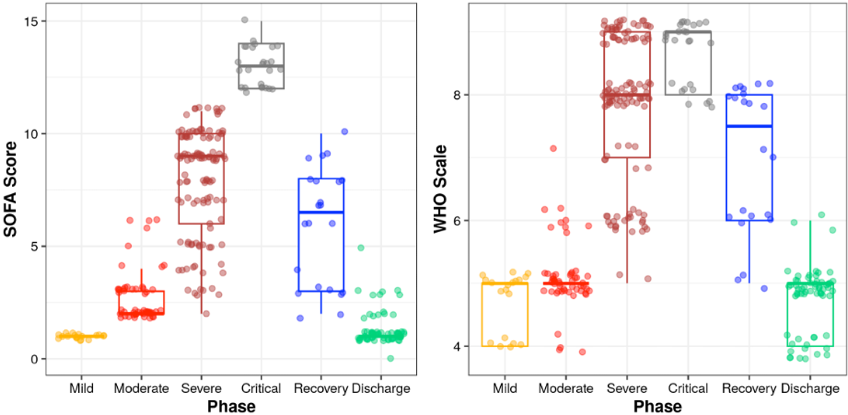

Figure S2. Relationship between phases and SOFA score and WHO scale.

As patients transition into more severe disease phases, SOFA and WHO scores increased, and as they recovered, SOFA and WHO scores decreased.

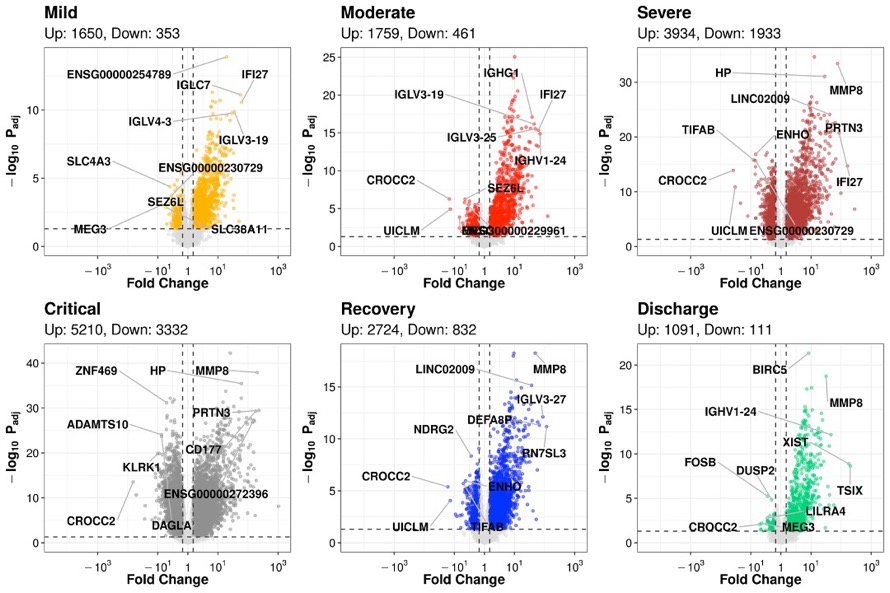

Figure S3. DE genes at each phase relative to follow-up controls increased with disease severity.

Coloured dots indicate DE genes (adj-p <0.05, absolute fold change ≥1.5). The top 5 up- and down- regulated genes (lowest adjusted p-value and highest fold change) are labelled.

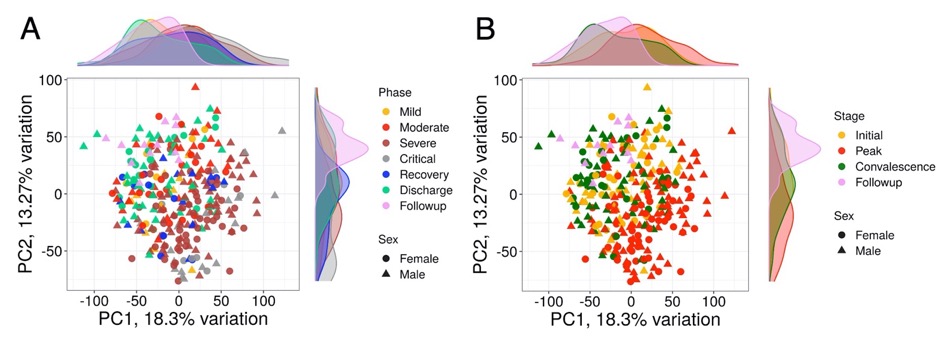

Figure S4. Samples separated based on disease phase and stage on PCA.

The first two principal components were plotted. Samples are coloured based on Phase (A) or Stage (B), and the shape corresponds to sex. Density plots on the sides show the distribution of samples based on Phase or Stage across the two principal components.

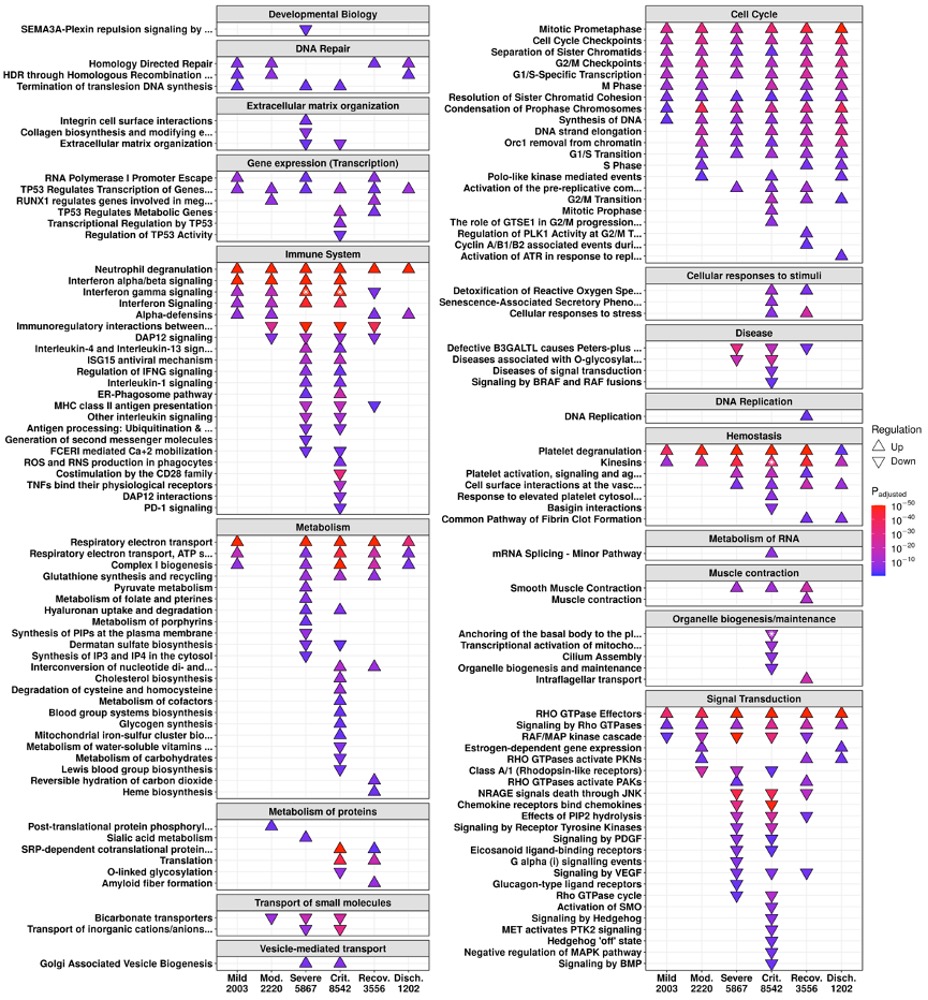

**Figure S5.** **All enriched Reactome pathways from DE genes of each phase compared to controls.** A subset of enriched pathways is displayed in **Figure 1A**. For certain pathways, both directions were enriched (indicated by *); the direction with the lower adjusted p-value (more significantly enriched) is shown. The total number of DE genes in each comparison are shown under each label. Mod: Moderate, Crit: Critical, Recov: Recovery, Disch: Discharge.

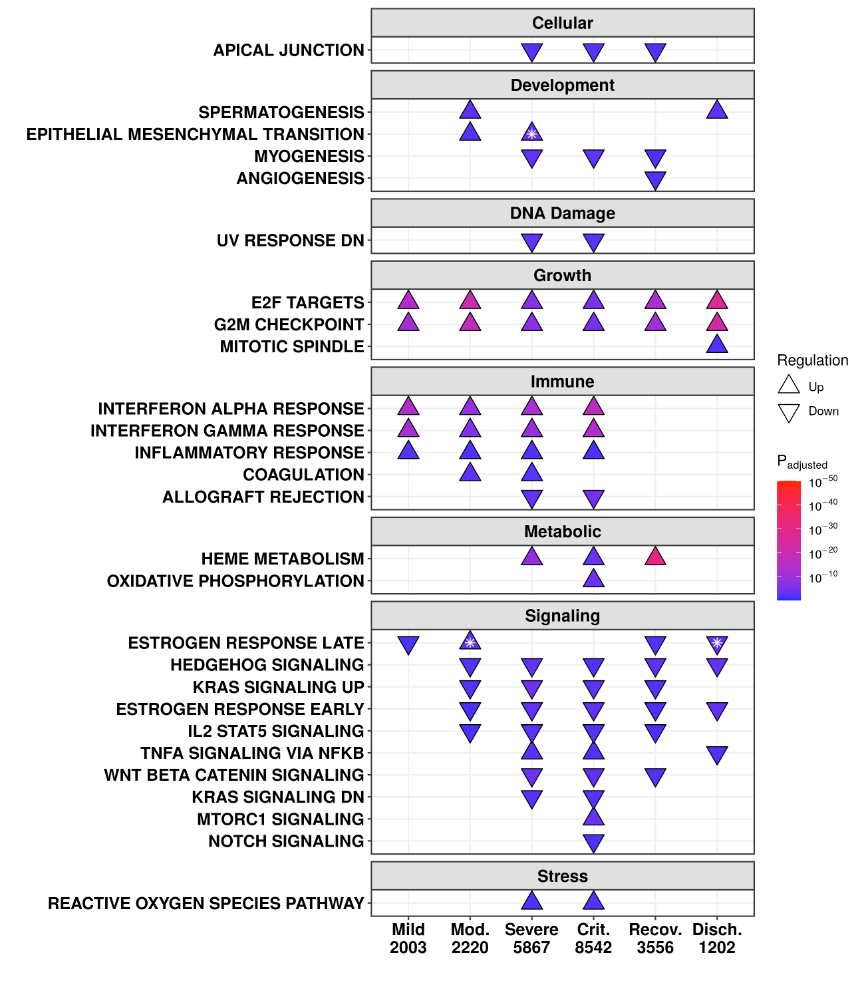

**Figure S6.** **All enriched Hallmark gene sets from DE genes of each phase compared to controls.** A subset of enriched gene sets is displayed in **Figure 1B**. For certain gene sets, both directions were enriched (indicated by *); the direction with the lower adjusted p-value (more significantly enriched) is shown. The total number of DE genes in each comparison are shown under each label. Mod: Moderate, Crit: Critical, Recov: Recovery, Disch: Discharge.

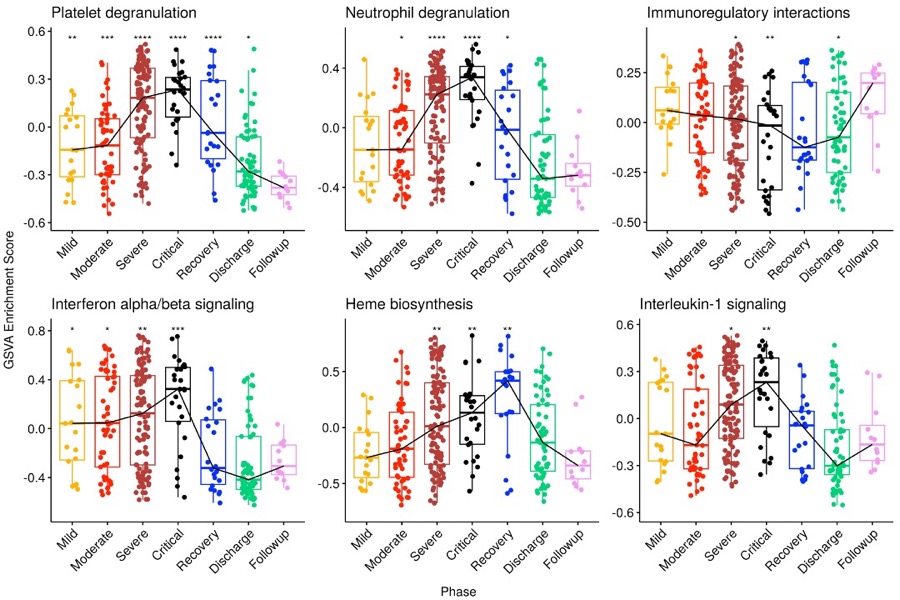

Figure S7. Temporal patterns of immune, hemostatic, and heme pathways vary throughout the disease timeline.

GSVA enrichment was performed using constituent genes of each pathway, with each point indicating the enrichment score of the pathway genes for each sample. The trend line connects median enrichment scores for each phase. A Wilcox test was performed for each phase relative to Follow-up (* = p<0.05, ** = p<0.01, *** = p<0.001, **** = p<0.0001).

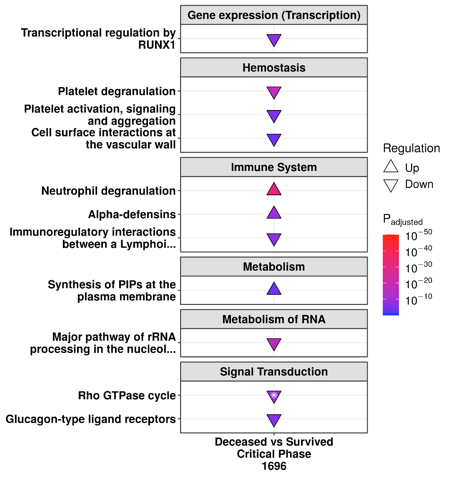

Figure S8. Aberrant immune and hemostasis pathways were seen in eventually deceased patients during the Critical phase compared to those who survived.

For one pathway, both directions were enriched (indicated by *); the direction with the lower adjusted p-value (more significantly enriched) is shown. The total number of DE genes in this comparison is shown under the label.

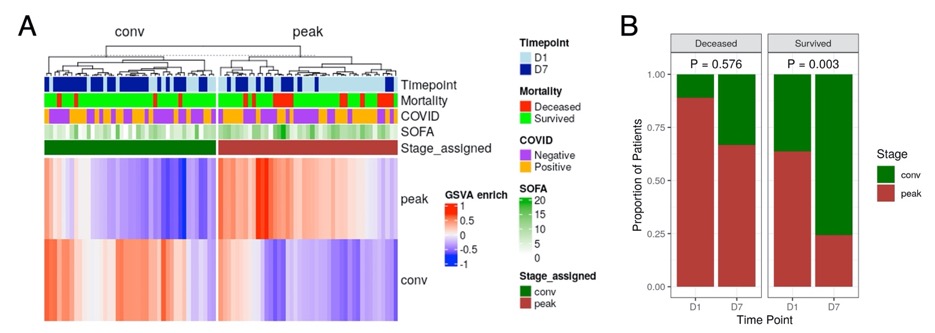

Figure S9. A significant number of survivors in the COLOBILI cohort transitioned from Peak to Convalescence stages during ICU hospitalization.

A: GSVA enrichment scores using condensed gene signatures of Peak and Convalescence stages for each sample. B: Proportion of non-survivors and survivors at each timepoint who were assigned to Peak or Convalescence stages. Chi-squared p-values are shown.
